# Unified rational protein engineering with sequence-only deep representation learning

**DOI:** 10.1101/589333

**Authors:** Ethan C. Alley, Grigory Khimulya, Surojit Biswas, Mohammed AlQuraishi, George M. Church

**Affiliations:** Wyss Institute for Biologically Inspired Engineering, Harvard University; Cambridge, MA 02138, USA; Department of Systems Biology, Harvard Medical School; Department of Genetics, Harvard Medical School

## Abstract

Rational protein engineering requires a holistic understanding of protein function. Here, we apply deep learning to unlabelled amino acid sequences to distill the fundamental features of a protein into a statistical *representation* that is semantically rich and structurally, evolutionarily, and biophysically grounded. We show that the simplest models built on top of this <u>uni</u>fied <u>rep</u>resentation (UniRep) are broadly applicable and generalize to unseen regions of sequence space. Our data-driven approach reaches near state-of-the-art or superior performance predicting stability of natural and *de novo* designed proteins as well as quantitative function of molecularly diverse mutants. UniRep further enables two orders of magnitude cost savings in a protein engineering task. We conclude UniRep is a versatile protein summary that can be applied across protein engineering informatics.

---

Protein engineering has the potential to transform synthetic biology, medicine, and nanotechnology. Traditional approaches to protein engineering rely on random variation and screening/selection without modelling the relationship between sequence and function^1, 2^. In contrast, rational engineering approaches seek to build quantitative models of protein properties, and use these models to more efficiently traverse the fitness landscape to overcome the challenges of directed evolution^3–9^. Such rational design requires a holistic and predictive understanding of structural stability and quantitative molecular function that has not been consolidated in a generalizable framework to date.

Although the set of engineering-relevant properties might be large, proteins share a smaller set of fundamental features that underpin their function. Current quantitative protein modeling approaches aim to approximate one or a small subset of them. For example, structural approaches, which include biophysical modeling^10^, statistical analysis of crystal structures^10^, and molecular dynamics simulations^11^, largely operate on the basis of free energy and thermostability in order to predict protein function. More data-driven co-evolutionary approaches rely on fundamental evolutionary properties to estimate the statistical likelihood of protein stability or function. While successful in their respective domains, these methods’ tailored nature, by way of the features they approximate, limit their universal application. Structural approaches are limited by the relative scarcity of structural data (Supp. Fig. 1), computational tractability, or difficulty with function-relevant spatio-temporal dynamics, which are particularly important for engineering^12–14^. Co-evolutionary methods operate poorly in underexplored regions of protein space (such as low-diversity viral proteins^15^) and are not suitable for *de novo* designs. Unlike these approaches, a method that scalably approximates a wider set of fundamental protein features could be deployed in a domain independent manner, bringing a more holistic understanding to bear on rational design.

**Figure 1.**
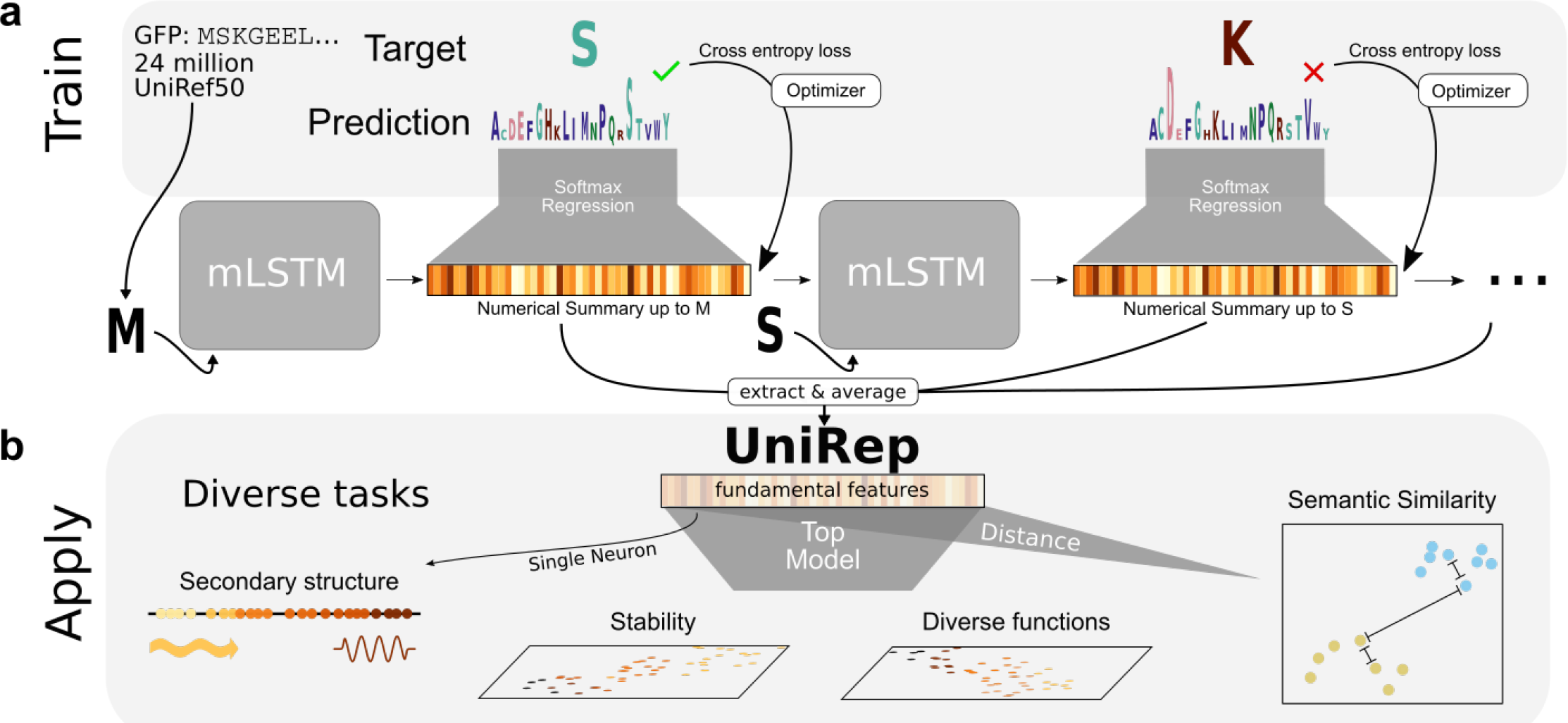
Workflow to learn and apply deep protein representations. **a.** UniRep model was trained on 24 million UniRef50 primary amino acid sequences. The model was trained to perform next amino acid prediction (minimizing cross-entropy loss), and in so doing, was forced to learn how to internally represent proteins. **b.** During application, the trained model is used to generate a single fixed-length vector representation of the input sequence by globally averaging intermediate mLSTM numerical summaries (the hidden states). A top model (e.g. a sparse linear regression or random forest) trained on top of the representation, which acts as a featurization of the input sequence, enables supervised learning on diverse protein informatics tasks.

Deep learning is a flexible machine learning paradigm that can learn rich data representations from raw inputs. Recently, this flexibility was demonstrated in protein structure prediction, replacing complex informatics pipelines with models that can predict structure directly from sequence^16^. Additionally, deep learning has shown success in sub-problems of protein informatics; for example: variant effect prediction^15^, function annotation^17, 18^, semantic search^18^, and model-guided protein engineering^3, 4^. While exciting advances, these methods are domain-specific or constrained by data scarcity due to the high cost of protein characterization.

On the other hand, there is a plethora of publicly available raw protein sequence data. The number of such sequences is growing exponentially^19^, leaving most of them uncharacterized (Supp. Fig. 1) and thus difficult to use in the modeling paradigms described above. Nevertheless, these are sequences from extant proteins that are putatively functional, and therefore may contain valuable information about stability, function, and other engineering-relevant properties. Indeed, previous works have attempted to learn raw sequence-based representations for subsequences^20^, and full-length “Doc2Vec” protein representations specifically for protein characteristic prediction^21^. However, these methods have neither been used to learn general representations at scale nor been evaluated on a comprehensive collection of protein informatics problems.

Here, we use a recurrent neural network to learn statistical representations of proteins from ∼24 million UniRef50^22^ sequences (Fig. 1a). Without structural or evolutionary data, this <u>uni</u>fied <u>rep</u>resentation (UniRep) summarizes arbitrary protein sequences into fixed-length vectors approximating fundamental protein features (Fig. 1b). This method scalably leverages underutilized raw sequences to alleviate the data scarcity constraining protein informatics to date, and achieves generalizable, superior performance in critical engineering tasks from stability, to function, to design.

## Results

### An mLSTM learns semantically rich representations from a massive sequence dataset

Multiplicative Long-Short-Term-Memory (mLSTM) Recurrent Neural Networks (RNNs) can learn rich representations for natural language, which enable state-of-the-art performance on critical tasks^23^. This architecture learns by going through a sequence of characters in order, trying to predict the next one based on the model’s dynamic internal “summary” of the sequence it has seen so far (its “hidden state”). During training, the model gradually revises the way it constructs its hidden state in order to maximize the accuracy of its predictions, resulting in a progressively better statistical summary, or *representation*, of the sequence.

We trained a 1900-hidden unit mLSTM with amino acid character embeddings on ∼24 million UniRef50 amino acid sequences for ∼3 weeks on 4 Nvidia K80 GPUs (Methods). To examine what it learned, we interrogated the model from the amino acid to the proteome level and examined its internal states.

We found that the amino-acid embeddings (Methods) learned by UniRep contained physicochemically meaningful clusters (Fig. 2a). A 2D t-Distributed Stochastic Neighbor Embedding^24^ (t-SNE) projection of average UniRep representations for 53 model organism proteomes (Supp. Table 1) showed meaningful organism clusters at different phylogenetic levels (Fig. 2b), and these organism relationships were maintained at the individual protein level (Supp. Fig. 2).

**Figure 2.**
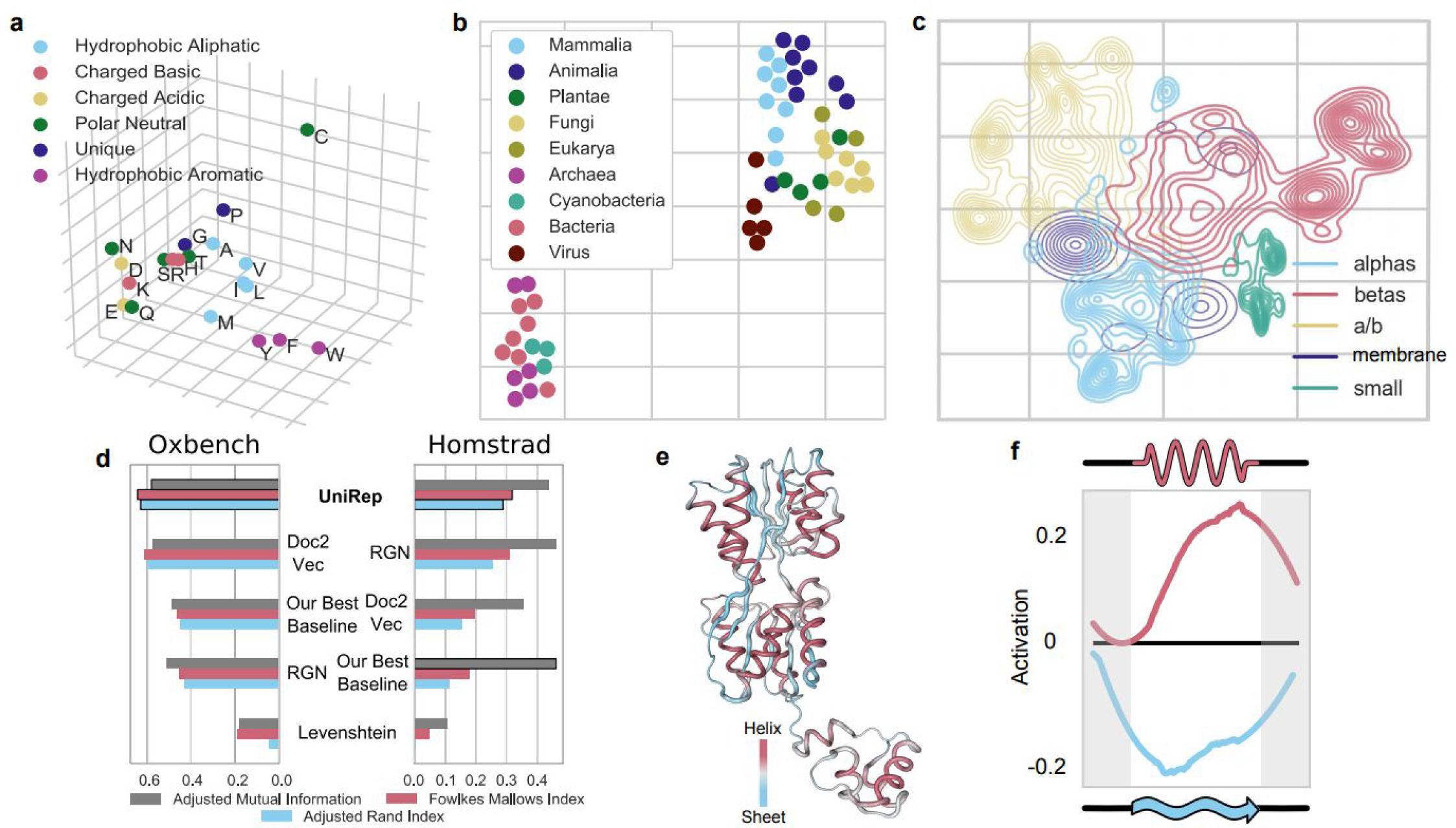
UniRep encodes amino acid physicochemistry, organism level information, secondary structure, evolutionary and functional information, and higher order structural features. **a.** PCA (Principal Component Analysis) of amino acid embeddings learned by UniRep. **b.** t-SNE of the proteome-average UniRep vector of 53 model organisms **c.** Low dimensional t-SNE visualization of UniRep represented sequences from SCOP colored by ground-truth structural classes, which were assigned after crystallization^29^. **d.** Agglomerative distance-based clustering of UniRep, a Doc2Vec representation method from Yang et al. (2018)^21^, a deep structural method from AlQuraishi (2018)^16^, Levenshtein (global sequence alignment) distance, and the best of a suite of machine learning baselines (Methods). Scores how well each approach reconstitutes expert-labeled family groupings from OXBench and HOMSTRAD. All metrics vary between 0 and 1, with 0 being a random assignment and 1 being a perfect clustering (Methods). **e.** Activation pattern of the helix-sheet secondary structure neuron colored on the structure of the Lac repressor LacI (PDB:2PE5, right). **f.** Average helix-sheet neuron (as visualized in f) activation as a function of relative position along a secondary structure unit (Methods).

To assess how semantically related proteins are represented by UniRep, we examined its ability to partition structurally similar sequences that share little sequence identity, and enable unsupervised clustering of homologous sequences.

UniRep separated proteins from various Structural Classification of Proteins (SCOP) classes derived from crystallographic data (Fig. 2c, Methods). More quantitatively, a simple Random Forest Model trained on UniRep could accurately group unseen proteins into SCOP superfamily and fold classes (Supp. Table 2, Methods).

We next sourced two expertly labeled datasets of protein families compiled on the basis of functional, evolutionary, and structural similarity: HOMSTRAD^25^ (3450 proteins in 1031 families) and OXBench^26^ (811 proteins in 180 families). Using Euclidean distances between UniRep vectors, we performed unsupervised hierarchical clustering of proteins in these families, and found good agreement with expert assignments according to three standard clustering metrics. We compared to baselines which include global sequence alignment distance computed with the Levenshtein algorithm, which is equivalent to the standard Needleman-Wunsch with equal penalties^27, 28^ (Fig. 2d, Supp. Figs. 3-5, Methods).

We finally examined correlations of the internal hidden states with protein secondary structure on a number of datasets (Methods). Surprisingly, we found a *single neuron* that discriminated beta sheets from alpha helices, positively correlating with alpha-helix annotations (Pearson’s *r*= .33, p < 1e-5), and negatively correlating with beta-sheet annotations (Pearson’s *r* = −0.35, p < 2e-6). Examination of its activation pattern on the Lac repressor structure visually confirmed these statistics (Fig. 2e). A larger-scale spatial analysis performed across many helices and sheets from different proteins revealed an activation pattern of the helix-sheet neuron that indicated it encodes features of both secondary structure units, going beyond individual amino acids (Fig. 2f). We found other neuron correlations with biophysical parameters including solvent accessibility (Supp. Fig. 6). Taken together, we conclude the UniRep vector space is semantically rich, and encodes structural, evolutionary, and functional information.

### UniRep enables stability prediction and generalizes to *de novo* designed proteins

Protein stability is a fundamental determinant of protein function and a critical engineering endpoint that affects the production yields^30^, reaction rates^31^, and shelf-life^32^ of protein catalysts, sensors, and therapeutics. Therefore, we next sought to evaluate whether UniRep could provide a suitable basis for stability prediction. Toward this end, we analyzed a large dataset of stability measurements for *de novo* designed mini proteins^5^. For proper model comparison we withheld a random test set never seen during training, even for model selection (Methods). We compared simple sparse linear models trained on top of UniRep (Fig. 1b) to those trained on a suite of baseline representations selected to encompass simple controls, standard models known to generalize well, and published state of the art. Among others, they included standard machine learning methods like bag-of-words, the state-of-the-art Doc2Vec representation from Yang *et al.* (2018)^21^, and a deep structural representation from the Recurrent Geometric Network (RGN)^16^ (Supp. Fig. 7, Methods). For this analysis, we also generated “UniRep Fusion” by concatenating the UniRep representation with other internal states of the mLSTM to obtain an expanded version of the representation (Methods, Supp. Table 3).

We also benchmarked against Rosetta, an established structural stability prediction method, using published Rosetta total energy scores for a subset of proteins in this dataset^5^. Despite lacking the explicit physical knowledge and structural data that Rosetta relies on, UniRep Fusion with a top model trained on experimental stability data significantly outperformed Rosetta on rank-order correlation with measured stability on the held-out test set (Spearman’s *ρ* = 0.59 vs. 0.42, Fig. 3a, Methods). Unlike UniRep, Rosetta does not provide a mechanism to incorporate our experimental stability data, which we recognize is a limitation of this comparison. UniRep Fusion additionally outperformed all baselines in our suite on the test dataset (Supp. Table 4-5, Methods). Due to Rosetta’s large computational requirements^33^ we did not extend this baseline to further analyses.

**Figure 3.**
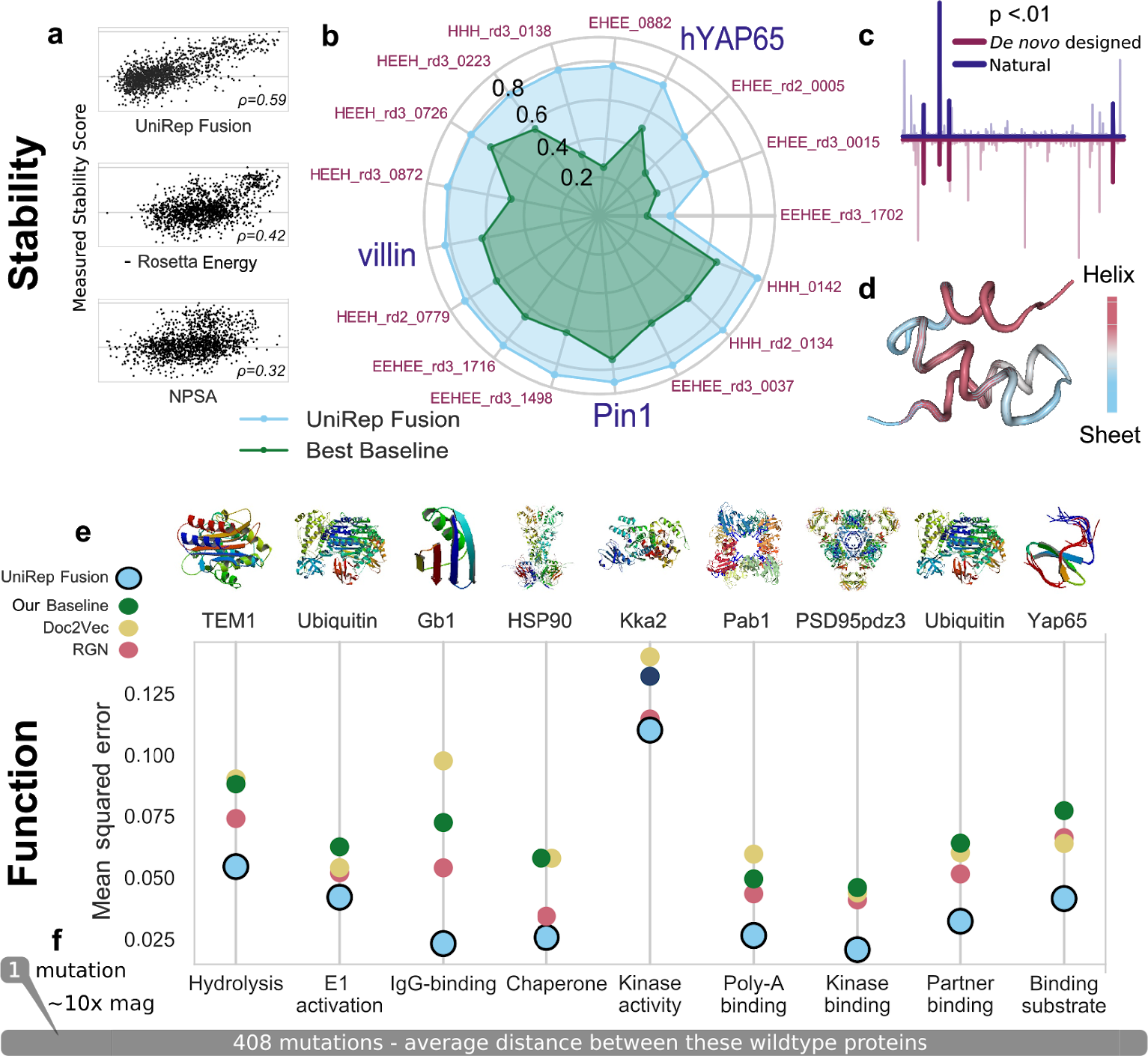
UniRep predicts structural and functional properties of proteins. **a.** Spearman correlation with true measured stability rankings of *de-novo* designed mini proteins UniRep Fusion-based model predictions and two alternative approaches: negative Rosetta total energy and non-polar surface area (Methods). UniRep Fusion outperforms both alternatives (*p* < 0.001; Welch’s t-test on bootstrap replicates). **b.** UniRep performance compared to a suite of baselines across 17 proteins in the DMS stability prediction task (Pearson’s *r*). UniRep Fusion achieved significantly higher Pearson’s *r* on all subsets (*p* < 0.006; Welch’s t-test on bootstrap replicates). **c.** Average magnitude neuron activations for *de novo* designed and natural protein stability prediction show significant co-activation (p < 0.01; permutation test). **d.** Activations of the helix-sheet neuron colored onto the *de novo* designed protein HHH_0142 (PDB: 5UOI). **e.** UniRep Fusion achieves statistically lower mean squared error than a suite of baselines across a set of 8 proteins with 9 diverse functions in the DMS function prediction task (p<0.0002 on 8/9 and p<0.009 on 1/9; Welch’s t-test on bootstrap replicates). **f**. UniRep performance across orders of scale: from distant proteins, to variants of the same protein one mutation apart. Scale bar illustrates the average distance between proteins in the DMS function prediction dataset. The scale of one mutation (10x magnification from the average distance bar) is shown to contrast the small mutational range UniRep is asked to model.

This result was surprising because *de novo* designed proteins constitute a miniscule proportion (1e-7) of the UniRep training data^34^. Thus, we further evaluated UniRep’s performance on *de novo* designs compared directly to natural proteins by using 17 distinct deep mutational scanning (DMS) datasets, which provide uniform measurements of stability of 3 natural and 14 *de novo* designed proteins^5^. These datasets consist of single-residue mutants, which presents an additional challenge for UniRep trained exclusively on sequences with <50% similarity. As before, a random test set was withheld for each protein.

We found UniRep Fusion-based models outperformed all baselines on both natural and *de novo* designed protein test sets. Surprisingly, the 3 proteins with the best test performance (EEHEE_rd3_0037, HHH_rd2_0134, HHH_0142 measured by Pearson’s *r*) were *de novo* designed (Fig. 3b, validation in Supp. Fig. 8). We confirmed these findings with a pooled analysis to train and predict on all subsets (Supp. Table 4-5). Further, we found that the same features of the representation were consistently identified by the linear model (Fig. 3c), suggesting a learned basis that is shared between *de novo* designed and naturally occurring proteins. Strikingly, the helix-sheet neuron, evaluated earlier on natural proteins (Fig. 2f), detected alpha helices on the *de novo* designed protein HHH_0142 (PDB: 5UOI) (Fig. 3d). Together, these results suggest that UniRep approximates a set of fundamental biophysical features shared between all proteins.

### UniRep enables prediction of the functional effects of single mutations for eight diverse proteins with distinct functions

Because UniRep enables prediction of stability, we hypothesized it could be a basis for the prediction of protein function directly from sequence. To test this hypothesis, we first sourced 9 diverse quantitative function prediction DMS datasets, incorporating data from 8 different proteins^35^. Each of these datasets only included single point mutants of the wild-type protein (>99% similarity), which were characterized with molecular assays specific to each protein function^35^. For each protein dataset, we asked if a simple sparse linear model trained on UniRep representations could predict the normalized (to wildtype) quantitative function of held out mutants.

On all 9 DMS datasets, UniRep Fusion-based models achieved superior test set performance, outperforming a comprehensive suite of baselines including a state-of-the-art Doc2Vec representation (Fig. 3e, Supp. Table 4-5). This is surprising given that these proteins share little sequence similarity (Fig. 3f), are derived from 6 different organisms, range in size (264 aa - 724 aa), vary from near-universal (hsp90) to organism-specific (Gb1), and take part in diverse biological processes (e.g. catalysis, DNA binding, molecular sensing, protein chaperoning)^35^. UniRep’s consistently superior performance, despite each protein’s unique biology and measurement assay, suggests UniRep is not only robust, but also must encompass features that underpin the function of all of these proteins.

### UniRep enables generalization through accurate approximation of the fitness landscape

A core challenge of rational protein engineering is building models which generalize from local data to distant regions of sequence space where more functional variants exist. Deep learning models often have difficulty generalizing outside of their training domain^36^. Unlike most models, which are only trained with data specific to the task at hand, UniRep was trained in an unsupervised manner on a wide sampling of proteins. Combined with its relative compactness (Supp. Fig. S9), we therefore hypothesized UniRep might capture general features of protein fitness landscapes which extend beyond task-specific training data.

To test this, we focused on fluorescent proteins, which have previously measured fitness landscapes^37^, and are useful for *in vivo* and *in situ* imaging, calcium and transmembrane voltage sensing, and optogenetic actuation^38^. We tested UniRep’s ability to accurately predict the phenotype of distant functional variants of *Aequorea victoria* green fluorescent protein (avGFP) given only local phenotype data from a narrow sampling of the avGFP fitness landscape^37^.

We considered what sized region of sequence space would make the best training data for UniRep. Training on a broad sequence corpus, like UniRef50, captures global determinants of protein function, but sacrifices fine-grained local context. On the other hand, training on a local region of extant sequences near the engineering target provides more nuanced data about the target, but neglects global features. Therefore, we hypothesized that an effective strategy may be to start with the globally trained UniRep and then fine-tune it to the evolutionary context of the task (Fig. 4a). To perform <u>evo</u>lutionary fine-<u>tuning</u> (which we call “evotuning”), we ran ∼13k unsupervised weight updates of the UniRep mLSTM (Evotuned UniRep) performing the same next-character prediction task on a set of ∼25k likely evolutionarily related sequences obtained via JackHMMER search (Methods). We compared this to the untuned, global UniRep as well as a randomly initialized UniRep architecture trained only on local evolutionary data (Evotuned Random; Fig. 4a, Methods).

**Figure 4.**
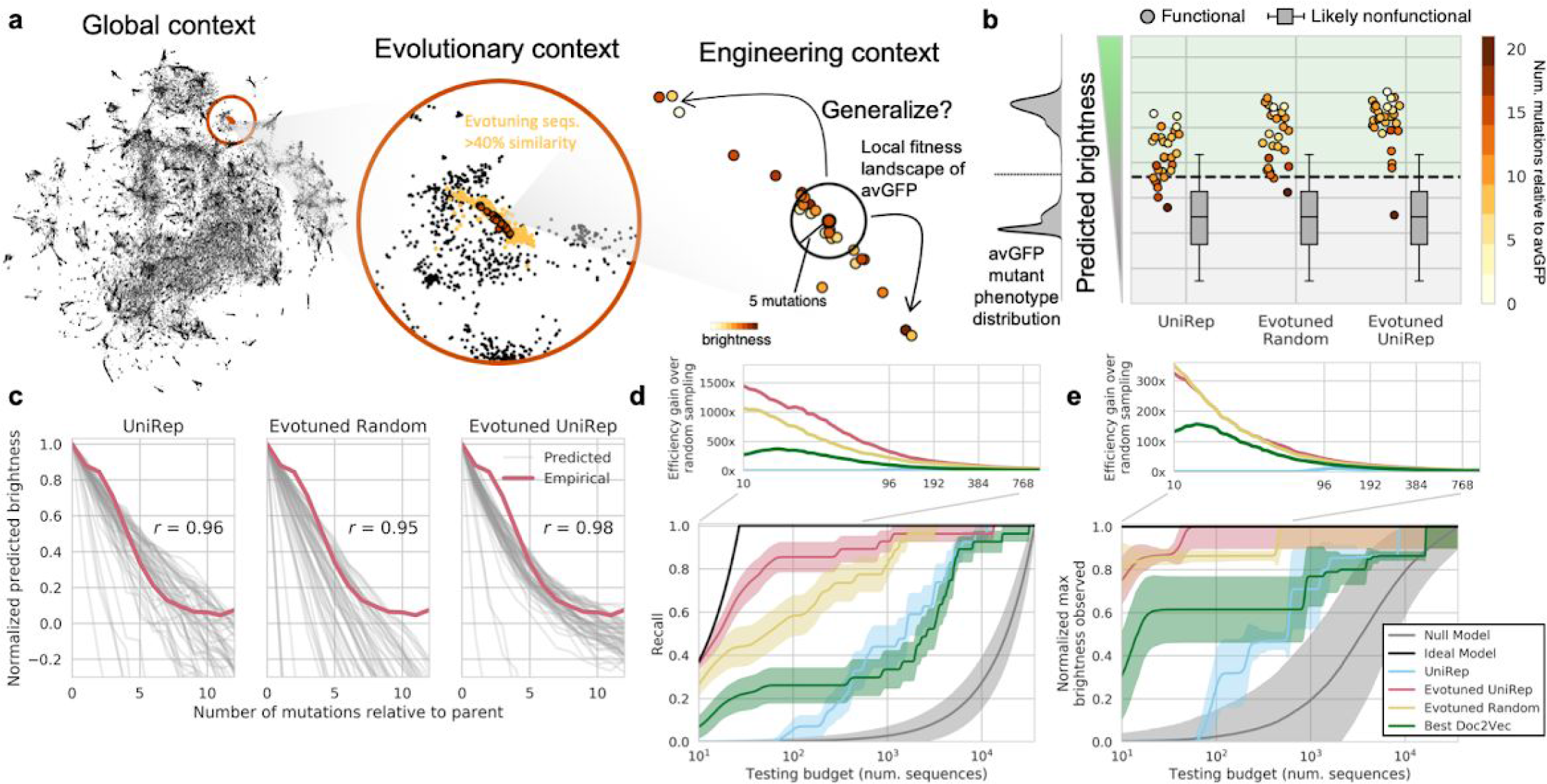
UniRep, fine-tuned to a local evolutionary context, facilitates protein engineering by enabling generalization to distant peaks in the sequence landscape. **a.** We hypothesize UniRep trades-off nuance for universality in a theoretical protein engineering task. By unsupervised training on a subspace of sequences related to the engineering target – “evotuning” – UniRep representations are honed to the task at hand. **b.** Predicted brightness of 27 homologs and engineered variants of avGFP under various representations + sparse linear regression models trained only on local avGFP data. Box and whisker plots indicate predicted distribution of dark negative controls. Green region above dotted line is predicted bright, below is predicted dark. On the left in gray is the training distribution from local mutants of avGFP. **c.** Predicted brightness-vs-mutation curves for each of the 27 avGFP homologs and engineered variants (the generalization set). Each grey line depicts the average predicted brightness of one of the 27 generalization set members as an increasing number of random mutations is introduced. Red line shows the average empirical brightness-vs-mutation curve for avGFP ^37^. **d.** Recall vs. sequence testing budget curves for each representation + sparse linear regression top model (bottom). Efficiency gain over random sampling (top) depicted as the ratio of a method’s recall divided by the recall of the null model as a function of testing budget. **e.** Maximum brightness observed vs. sequence testing budget curves for each representation + sparse linear regression top model (bottom). Efficiency gain over random sampling analogously defined for recall but instead with normalized maximum brightness (top).

Using these trained unsupervised models we generated representations for the avGFP variant sequences from Sarkisyan *et al.* (2016)^37^ and trained simple sparse linear regression top models on each to predict avGFP brightness. We then predicted the brightness of 27 functional homologs and engineered variants of avGFP sourced from the FPbase database^39^ using each of the models. The sequences in this generalization set were 2 to 19 mutations removed from avGFP, and were not present in the set of sequences used for evotuning.

All three representation-based models correctly predicted most of the sequences in the generalization set to be bright (Fig. 4b), with the Evotuned UniRep based model having the best classification accuracy (26/27 correctly classified). As a control, we predicted the brightness of 64,800 variants, each of which harbored 1-12 mutations relative to a generalization set member. We assumed variants with 6-12 mutations were non-functional based on empirical observation of the avGFP fitness landscape^37^. Indeed, variants with 6-12 mutations were predicted to be non-functional on average, confirming that these representation + avGFP top models were both sensitive and specific even on a distant test set (Fig. 4b). Figure 4c further illustrates that, on average, the representation + avGFP top models predicted decreasing brightness under increasing mutational burden. In particular, the Evotuned UniRep-based predictions for generalization set members closely matched the empirical brightness-vs-mutation curve for that of avGFP^37^ (Pearson’s r=.98). These observations imply that each model predicts generalization set members sit at a local optima in the fitness landscape. Furthermore, Evotuned UniRep, unlike the Evotuned Random, predicted a non-linear decline in brightness indicative of predicted epistasis, which is consistent with previous theoretical and empirical work on epistasis in protein fitness landscapes^37, 40, 41^. These data suggest that UniRep-based models generalize by building good approximations of the distant fitness landscape using local measurements alone.

We confirmed these findings with additional analysis of generalization for distinct prediction tasks (Supp. Fig. 10). We show that UniRep does not require evotuning to enable generalizable prediction of stability (Supp. Fig 10c), even if forced to extrapolate from nearby proteins to more distant ones (Supp. Fig 10d). We also provide data from a variant effect prediction task, which enumerates the requirements – such as standardized phenotype measurement – of an appropriate generalization task for a sequence-only model like UniRep (Supp. Fig. 10b).

To better understand how UniRep enables extrapolation, we visually examined the spatial pattern of training and test data in sequence space (Supp. Fig 10e), Doc2Vec space (Supp. Fig 10g), and UniRep space (Supp. Fig. 10f,h). Interestingly, despite not overlapping in sequence space, training and test points were co-localized in UniRep space suggesting that UniRep discovered commonalities between training and test proteins that effectively transformed the problem of extrapolation into one of interpolation, providing a plausible mechanism for our performance.

### Evotuning of UniRep increases efficiency and decreases cost in a fluorescent protein engineering problem

We hypothesized that UniRep’s generalizable function prediction could enable the discovery of functional diversity and function optimization — the ultimate goals of any protein engineering effort. We constructed a sequence prioritization problem in which the representation + avGFP top models were tasked with prioritizing the 27 truly bright homologs from the generalization set over the 32,400 likely non-functional sequences containing 6-12 mutations relative to a member of the generalization set. We tested each model’s ability to prioritize the brightest GFP in the dataset under differently sized testing budgets, and its ability to recover all bright members with high purity. “Brightest sequence recovery” measures a model’s utility for function optimization. “Bright diversity recovery,” as measured by statistical recall, measures the ability of each model to capture functional diversity, which could represent engineering endpoints or substrates for directed evolution. As a lower-bound model, we defined a null model that orders sequences randomly, and as a upper-bound model, we defined an ideal model that prioritizes sequences perfectly according to ground truth. As a baseline, we trained avGFP top models on Doc2Vec representations^21^ (Methods).

Figures 4d,e illustrate each representation + avGFP top model performance in terms of recall and prioritization of sequences by brightness. All models outperformed the null model in terms of recall and brightest sequence recovery. While the best performing Doc2Vec baseline outperformed untuned UniRep in both tasks, the various Doc2Vec baselines were unreliable in performance (Supp. Fig. 11) in a manner that was not explainable by their expressivity, architecture, or training paradigm. Evotuned UniRep demonstrated superior performance in both the function diversity and function optimization tasks, having near ideal recall for small (<30 sequence) testing budgets and quickly recovering the brightest sequence in the generalization set of >33,000 sequences with fewer than 100 sequences tested (Fig. 4d, red). The global information captured by UniRep was critical for this gain as evotuning from a random initialization of the same architecture yielded inferior performance (Fig. 4d, yellow). More concretely, for small plate-scale testing budgets of 10-96 sequences, Evotuned Unirep achieved a ∼3-5x improvement in recall and a ∼1.2-2.2x improvement in maximum brightness captured over the best Doc2Vec baseline at the same budget. Conditioning on a desired performance level, Evotuned UniRep achieved 80% recall within the first ∼60 sequences tested, which would cost approximately $3,000 to synthesize. By contrast, the next best approach examined here would be 100x more expensive (Supp. Fig. 12).

## Discussion

In this work we used abundant unlabeled protein sequence data to learn a broadly applicable statistical vector representation of proteins. This representation enabled near state-of-the-art or superior performance on more than 20 stability and function prediction tasks (Supp. Table 6) that reflect the challenges faced by protein engineers. Importantly, these results (Fig. 3, Supp. Table 4-5) were obtained using the same set of UniRep-parameterized features, which were superior to a suite of non-trivial baseline representations. Given additionally the simplicity of the top-models used and the compactness of UniRep (Supp. Fig. 9), this provides strong, albeit indirect, evidence that UniRep must approximate fundamental protein features that underpin stability and a broad array of protein-specific functions.

A more direct interrogation of UniRep features at the amino acid to whole proteome level revealed that these features at least embody a subset of known characteristics of proteins (Fig. 2); however, we note the possibility of UniRep representing more. Because UniRep is learned from raw data, it is unconstrained by existing mental models for understanding proteins, and may therefore approximate currently unknown engineering-relevant features. Taken together, these results suggest UniRep is likely to be a rich and robust basis for protein engineering prediction tasks beyond those examined here.

Remarkably, UniRep achieves these results from “scratch”, using only raw sequences as training information. This is done without experimentally determining, or computationally folding, a structural intermediate - a necessary input for alternative methods^6, 42^. By enabling rapid generalization to distant, unseen regions of the fitness landscape, UniRep may improve protein engineering workflows, or in the best case, enable the discovery of sequence variants inaccessible to purely experimental or structural approaches. Although the utility of the representation is limited by sampling biases in the sequence data^43, 44^ length of training, the size^45^ and coverage^43^ of sequence databases as well as deep-learning specific computational hardware^46^ are improving exponentially. Coupled with the continued proliferation of cheap DNA synthesis/ assembly technologies^47^, and methods for digitized and multiplexed phenotyping, UniRep-guided protein design promises to accelerate the pace with which we can build biosensors^13^, protein^48^ and DNA binders^49^, and genome editing enzymes^50^.

Finally, there are many exciting, natural extensions of UniRep. It can already be used generatively (Supp. Fig. 13), evoking deep protein design, similar to previous work with small molecules^51^. Beyond engineering, our results (Fig. 2b-e, Supp. Fig. 2) suggest UniRep distance might facilitate vector-parallelized semantic protein comparisons at any evolutionary depth. Among many straightforward data augmentations (Supp. Table. 7), UniRep might advance ab-initio structure prediction by incorporating untapped sequence information with the RGN^52^ via joint training^52^. Most importantly, UniRep provides a new perspective on these and other established problems. By learning a new basis directly from ground-truth sequences, UniRep challenges protein informatics to go directly from sequence to design.

## Acknowledgements

We thank John Aach, Amaro Taylor-Weiner, Daniel Goodman, Pierce Ogden, Gleb Kuznetsov, Sam Sinai, Aaron Tucker, Miles Turpin, Jacob Swett, Nathaniel Thomas, Raahil Sha, Chris Bakerlee and Kyle Fish for valuable feedback and discussion. S.B. was supported by an NIH Training Grant to the Harvard Bioinformatics and Integrative Genomics program as well as an NSF GRFP Fellowship. M.A. was supported through NIGMS Grant P50GM107618 and NIH grant U54-CA225088. E.C.A. and G.K. were supported by the Center for Effective Altruism. E.C.A. was partially supported by the Wyss Institute for Biologically Inspired Engineering. Computational resources were, in part, generously provided by the AWS Cloud Credits for Research program.

## Author Contributions

E.C.A. and G.K. conceived the study. E.C.A., G.K., and S.B. conceived the experiments, managed data, and performed the analysis. M.A. performed performed large-scale RGN inference and managed data and software for parts of the analysis. G.M.C. supervised the project. E.C.A., G.K. and S.B. wrote the manuscript with help from all authors.

## Competing Interests

E.C.A., G.K., and S.B. are in the process of pursuing a patent on this technology. S.B. is a former consultant for Flagship Pioneering company VL57. A full list of G.M.C.’s tech transfer, advisory roles, and funding sources can be found on the lab’s website: http://arep.med.harvard.edu/gmc/tech.html

## Code Availability

Code for UniRep model training and inference with trained weights along with links to all necessary data is available at https://github.com/churchlab/UniRep.

## Data Availability

All data are available in the main text or the supplementary materials.

## Materials and Methods

### Training the UniRep representations

#### Unsupervised training dataset

We expected that public protein databases, unlike many Natural Language datasets, would contain a) random deleterious mutations yet to be eliminated by selection, and b) hard-to-catch sequencing/assembly mistakes, both leading to increased noise. Therefore, we chose UniRef50^22^ as a training dataset. It is “dehomologized” such that any two sequences have at most 50% identity with each other. By selecting the single highest quality sequence for each cluster of homologs^22^, we hypothesized UniRef50 would be less noisy. It contains ∼27 million protein sequences. We removed proteins longer than 2000 amino acids and records containing non-canonical amino acid symbols (X, B, Z, J), randomly selected test and validation subsets for monitoring training (1% of the overall dataset each) and used the rest of the data (∼24 million sequences) in training.

#### Models and training details

We approached representation learning for proteins via Recurrent Neural Networks (RNNs). Unlike other approaches to representing proteins, namely as one-hot-encoded matrices as in Biswas et. al 2018 ^3^, RNNs produce fixed-length representations for arbitrary-length proteins by extracting the hidden state passed forward along a sequence. While padding to the maximum sequence length can in principle mitigate the problem of variable length sequences in a one hot encoding, it is ad-hoc, can add artifacts to training, wastes computation processing padding characters, and provides no additional information to a top model besides the naive sequence. Furthermore, even very large representations, like the 1900 dimensional UniRep, are more compact than average protein length 1-hot encodings (Supp. Fig. 9), reducing the potential for overfitting. While Doc2Vec methods produce fixed-length vectors, they have been empirically outperformed by more expressive architectures like RNNs in recent work on representation learning in natural language ^23^.

We considered the mLSTM^53^, LSTM^54^, and Gated Recurrent Unit (GRU)^55^ for candidate RNN representation learners. After manual explorations comparing these classes, and considering previous work demonstrating the success of the mLSTM for a similar task in natural language^23^, we decided to use the mLSTM. Specifically, the architectures selected for large-scale training runs were a 1900-dimensional single layer multiplicative LSTM^53^ (∼18.2 million parameters) as described elsewhere^23^, a 4-layer stacked mLSTM of 256 dimensions per layer (∼1.8 million parameters), and a 4-layer stacked mLSTM with 64 dimensions per layer (∼.15 million parameters), all regularized with weight normalization^56^. As a matter of definition, we note that because all of these networks are recurrent, even the single hidden layer mLSTM-1900 is considered “deep” because the network is unrolled in the timestep dimension as a composition of hidden layers through time.

We followed a heuristic that assumes, for large data sets like ours, more expressive models will learn richer representations. Thus, we selected 1900 dimensions in the large single-layer mLSTM because it was approximately the largest dimensionality that could fit in GPU memory after some experimentation. We tried smaller widths (the 256 and 64 dimensions) in case the large number of latent dimensions in the 1900-unit mLSTM led to overfitting on prediction tasks. Our comparison with the mLSTM-1900 suggested this was almost never the case (Supp. Data 3).

Sequences of amino acids were one-hot encoded and passed through a 10 dimensional amino-acid character embedding before being input to the mLSTM layer. For the smaller stacked networks, both standard recurrent and residual recurrent connections, in which the hidden states of each layer are summed, were evaluated. For these stacked networks, dropout probability was selected from {0, .5}. Hyperparameters were tuned manually on a small number of weight updates and final parameters were selected based on the rate and stability of generalization loss decrease. We found that dropout and residual connections both increased validation set error. We hypothesized that residual connections, which should improve gradient flow to earlier layers, were not advantageous here given the small number of layers tested. We further hypothesized that these networks did not require dropout or other regularization outside of weight normalization because of the high ratio of observations to model parameters.

All models were trained with the Adam optimizer using truncated-backpropagation through time with initial states initialized to zero at the beginning of sequences and persistent across updates to simulate full backpropagation as described previously^23^. Batch sizes and truncation windows were selected to fit into GPU memory and were, respectively, 256 and 128 (mLSTM-1900), 512 and 384 (4x-mLSTM-256), 1028 and 384 (4x-mLSTM-64). Training was performed using data parallelism on 4 Nvidia K-80 GPUs (mLSTM-1900) or 2 Nvidia K-40s (4x-mLSTM-256, 4x-mLSTM-64). The mLSTM-1900 model was trained for ∼770K weight updates, or ∼3.5 weeks wall clock time, corresponding to ∼1 epoch. The 4x-mLSTM-256 and 4x-mLSTM-64 were trained for ∼90K weight updates, ∼2 days wall clock time, ∼3 epochs and 220k weight updates, ∼2 days wall clock time, 14 epochs, respectively.

#### Computing vector representations

The mLSTM architecture has two internal states that encode information about the sequence it is processing, the hidden state and the cell state^53^. One hidden state and one cell state are produced for every amino acid in a forward pass over a sequence. Previous work in Natural Language has used the final hidden state (corresponding to the residue at the C terminus in our case) as the sequence representation^23^.

Compared to natural language, we hypothesized that the complexity of protein folding would generate more long-range and higher-order dependencies between amino acids. Therefore, we elected to construct the UniRep representation as the *average* of the 1900-unit model’s hidden states, integrating information across distant amino-acids. We hoped this would better represent long-term dependencies critical to protein function prediction.

For convenience we named this vector representation – the Average Hidden state of mLSTM-1900 – simply “UniRep” everywhere in the main text.

We compared the performance of UniRep as defined above with other possibilities for the representation state. We extracted final hidden state produced by the model when predicting the last amino acid in a protein sequence (Final Hidden) and the last internal cell state (Final Cell).

Curious whether these vectors contained complementary information, we also constructed a concatenation of all 3 representation possibilities (Average Hidden, Final Hidden and Final Cell) from the 1900-unit mLSTM. For convenience, we named this 1900 x 3 dimensional representation “UniRep Fusion” everywhere in the main text. We built various other concatenations of these representations (Supp. Table 3), denoted as “Fusions”, to be evaluated in supervised stability and quantitative function prediction.

We consider UniRep to be the most information-dense representation evaluated here, while UniRep Fusion is the most complete. UniRep, by virtue of its smaller dimensionality, should be deployed where computational resources are constrained. We present UniRep Fusion only for supervised prediction tasks (of protein stability, biophysical characteristics and function).

For completeness we evaluated the influence of network size in representation performance. There is evidence that larger neural networks learn better representations^23^. When we compared UniRep, UniRep Fusion and the other 1900-unit representations defined above to identical representations extracted from 64 and 256-unit models with an identical architecture (training described above, Supp. Table 3) our results agree with this pattern except with very small datasets, which are more variable as expected (Supp. Data 1,3).

### Supervised model benchmarking

#### Sourcing and processing analysis datasets

All the datasets (Supp. Table 6) were obtained from the supplemental information of corresponding publications. Sequence, function pairs were deduplicated, validated to exclude records containing non-standard amino acid symbols, and split randomly into 80-10-10 train - validation - test sets as described below. Size is reported after cleaning. When protein sequences could not be found in the published data, they were retrieved from UniProt by whatever sequence identifiers were available, using UniProt ID mapping utility (https://www.uniprot.org/help/uploadlists). After this, the same split and top model training analysis was done with all datasets, as described below.

Inference of RGN representations on a small number (on the order of 10s) of sequences could not be completed due to challenges constructing PSSMs using the JackHMMER program (the multiple sequence alignments were so large they would not fit in available memory). To obtain comparable scores between the performance of RGN and other representations, we dropped these sequences from the datasets used for training and evaluation.

#### Train-Validation-Test Split

For most supervised tasks no test set was provided, so we made a 80%-10%-10% Train-Validation-Test split in python using the numpy package and a fixed random seed. The validation set was never used for training so that it could be used to estimate generalization performance while we were conducting experiments and building models. In order to obtain a final, truly held-out, measure of generalization performance we did not look at the test set until all models were finalized and a submission ready version of this manuscript was prepared.

#### Baselines representations - RGN

We used the Recurrent Geometric Network (RGN) model trained on the ProteinNet12 dataset^57^ and assessed on CASP12 structures in AlQuraishi (2018)^16^. The model was used as is without additional training or fine-tuning. Instead of using the predicted structures of the RGN, we extracted the internal state learned by the model for each sequence + PSSM combination (3200 dimensions corresponding to the last outputs of two bidirectional LSTMs, each comprised of 800 units per direction). PSSMs were generated in a manner identical to that used for generating ProteinNet12 PSSMs^57^.

#### Baseline representations - Doc-2-Vec

We used the 4 best performing models as chosen for presentation by the authors of Yang *et al.* (2018)^21^. These are (using the original names): original with k=3 w=7, scrambled with k=3 w=5, random with k=3 w=7, and uniform k=4 w=1. We downloaded the trained models from http://cheme.caltech.edu/~kkyang/models/ and performed inference on all of the sequences in our datasets, adapting as much as possible the code used by the authors found at https://github.com/fhalab/embeddings_reproduction.

#### Baseline representations - N-gram, length, and basic biophysical

We utilized 6 baseline representations - they constitute 3 distinct approaches to constructing a general scalable protein vector representations without using deep learning techniques:

- Amino-acid frequencies in the protein and protein length normalized to average protein length in the unsupervised training data
- Amino-acid frequencies in the protein concatenated with predicted biophysical parameters of the protein: molecular weight, instability index, isoelectric point, secondary structure fraction. The annotation/prediction for biophysical parameters was done with biopython package version 1.72 (https://biopython.org/)
- Two n-gram representations with n=2 and n=3 (with scikit-learn 0.19.2 http://scikit-learn.org/stable/index.html).
- Two n-gram representations with n=2 and n=3 with Term Frequency - Inverse Document Frequency (TF-IDF) weighting intended to emphasize the n-grams unique to particular proteins^58^

See Supp. Table 8 for an exhaustive list of all the baseline variants used.

#### Stability Ranking Task

To further benchmark our performance compared to typical protein engineering tools, we obtained the published Rosetta total energy estimates with the “beta_nov15” version of the energy function^59, 60^ and exposed nonpolar surface area of the designed structure for the stability prediction for *de novo* designed mini proteins dataset^5^. The Rosetta calculations were not performed in^5^ for the control proteins in this dataset, so the scores were available for 1432 out of 5570 test set proteins and 1416 out of 5571 validation set proteins in our splits. Because these Rosetta Total Energy estimates do not directly correspond to measured stability values (e.g. lower energy is higher stability as defined), we used Spearman Rank Order Correlation coefficient, comparing our model predictions and the alternative scores’ ability to reconstitute correct stability ranking of the sequences in the dataset.

#### Regression analysis with Lasso LARS

We utilized a simple sparsifying linear model with L1 prior using the Least Angle Regression (LARS) algorithm (implementation from the scikit-learn package 0.19.2 http://scikit-learn.org/stable/index.html). The value of regularization parameter alpha for the model for each representation on each task was selected through 10-fold random cross-validation on the training data.

We obtained an estimate of the standard deviation of the resulting validation/test metrics by resampling 50% of the validation/test set 30 times and computing an empirical standard deviation of the obtained metrics, and used Welch’s t-test for comparisons.

We were unable to compare to the published state of the art on the 9 DMS function prediction datasets from Gray *et al.* (2017) ^35^. The exploratory nature of the authors’ analysis of these datasets for in-domain prediction led them to use their test set during hyperparameter selection, making their results incomparable with the best machine learning practices we followed here. This is in contrast to out-of-domain/transfer performance, which was the main objective of the Gray et al’s (2017) analysis. Unlike in-domain prediction, Gray et al (2017) followed best practices for leave one protein out (LOPO) transfer analysis (see below).

### Supplemental generalization analysis

#### Leave-one-protein-out transfer

We evaluated generalization (transfer) learning performance using the standard procedure based on previous foundational work, Glorot *et al.* (2011)^61^. This particular analysis was done for the quantitative function prediction (including data for 8 different proteins) and for 17 protein DMS stability prediction datasets.

Briefly, e(S, T) - the *transfer error* - is defined as the test/validation error achieved by a model trained on the source domain S and evaluated on the target domain T. e(T, T) is similarly defined and is called the *in-domain error*.

In the case of quantitative function prediction and stability prediction (natural and *de novo*), we used leave-one-protein out approach, constructing a Source/Target split for each protein in the dataset with that protein as the Target and the rest of the proteins as the Source. We used the same linear model we used in in-distribution regression analysis (L1 prior, LARS algorithm). The value of regularization parameter alpha was in turn obtained through leave-one-protein-out cross-validation on the training data separately for each split.

To control for the differences in difficulty between various splits, we also evaluated the *baseline in-domain error* - eb(T, T) - the error achieved by a baseline representation when trained and evaluated on the Target domain. We used amino acid frequency and protein length as our baseline representation, and computed eb(T, for each of the target domains defined above.

Transfer ratio is the ratio of transfer error to baseline in-domain error e(S, T)/eb(T, T). We use the average of transfer ratios over all Source/Target splits for a given representation to characterize transfer performance on the given dataset. We also look at the in-domain ratio e(T, T)/eb(T, T), which reflects the in-distribution performance in comparable terms to the transfer ratio.

We obtained an estimate of the standard deviation of the resulting validation/test metrics by computing an empirical standard deviation of metrics across all the different hold-one-out splits.

In the case of 17 DMS stability datasets (Supp. Table 6), we also constructed an additional extrapolation Source/Target split - from central to remote proteins (Supp. Fig. 10d) as follows. We computed a string median of initial sequences of 17 proteins in these datasets, and selected the 4 proteins with the largest edit distance from the median. We then computed a multidimensional scaling (MDS) 2D plot of the Levenshtein distance matrix of the 17 initial proteins, and selected 4 most peripheral proteins along each axis of the plot (Supp. Fig. 10e). Together, the DMS datasets for these 8 proteins constituted the Target dataset (also known as test set shown in Red on Supp. Fig. 10e) to evaluate our generalization/transfer performance. The other 9 datasets served as the Source dataset to be trained on. Once the Source/Target split was defined, the transfer analysis was performed similarly to the above except that we did not average any transfer metrics (because in this case there is only a single Source and Target).

### Supervised remote homology detection

#### Datasets - Håndstad fold and superfamily

We used two standard benchmarks based on the SCOP database from Håndstad et al.^62^: the superfamily-level remote homology detection and the harder fold-level similarity detection.

Briefly, the superfamily benchmark sets up a binary classification problem for each of the superfamilies in the dataset: a single family in the superfamily presenting a positive test set, the other families in that superfamily serving as the positive training set, a negative test set comprising one random family from each of the other superfamilies, and the negative training set combining the rest of the families in these superfamilies.

The fold-level benchmark is analogous at the fold level, setting up a classification problem for each of the folds in the dataset: one superfamily in the fold is used as positive test set, the others in that fold serving as the positive training set, a single random superfamily from every other fold comprising a negative test set, and taking the remaining sequences as the negative training set.

This structured training/ test set could not be straightforwardly subsampled, so to protect against overfitting we instead held out training and evaluation on the entire fold-level dataset until the model was trained and our Bayesian hyperparameter tuning procedure finalized on the superfamily-level benchmark dataset. Because these are standard benchmarking datasets and the task was computationally expensive, we choose to evaluate only UniRep performance. Many of the more recent published methods for remote homology detection ^63, 64^ use PSSMs as a source of evolutionary information, which we excluded for equal comparison to UniRep, which was trained on strictly dehomologized sequences and had no access to local evolutionary information like a PSSM.

#### Binary classification with Random Forests

For supervised remote homology benchmarks we used Random Forest implementation from the same scikit-learn package with 1000 estimators and a “balanced” class weight. For each of the binary classification tasks in each benchmark, we conducted a Bayesian hyperparameter optimization as implemented in the skopt package (https://scikit-optimize.github.io/), picking the following hyperparameters: maximum proportion of features to look at while searching for the best split (between 0.5 and 1), function to measure the quality of a split (Gini impurity or information gain), the minimum number of samples required to be at a leaf node (between 0.1 and 0.3), and the minimum number of samples required to split an internal node (between 0.01 and 1) with 75 iterations of optimization, 3-fold cross-validation on the training data and ROC score as the scoring metric.

To make our model comparable with the scores of previously developed remote homology detection models in the literature, we used two standard metrics - ROC score (normalized area under the receiver operating characteristic curve) and ROC50 (ROC score at the point where the first 50 false positives occur).

Two out of one hundred and two superfamilies, identified in the data through the name of the family presenting the positive test set - c.2.1.3 and c.3.1.2, and one out of eighty six folds, identified in the data by the name of the superfamily representing the positive test set - c.23.12, were excluded from the consideration due to our failure to obtain a coherent positive/negative train/test split from the published data source^64^.

### Unsupervised clustering of distant but functionally related proteins

#### Agglomerative clustering of representations and hierarchical clustering confirmation

We used basic agglomerative clustering (implemented in python with sklearn^65^) with the average linkage metric and Euclidean distance for all vector representations, and a precomputed Levenshtein distance matrix using the python-Levenshtein package for the sequence-only control.

To confirm our cluster assignments, we picked a small set of 3 Cytochrome Oxidase (1,2,3) families from OXBench, 8 proteins total. We used the fastcluster package to produce a linkage matrix with average linkage and Euclidean distance (for UniRep) and average linkage and Levenshtein distance, implemented in python-Levenshtein, for the sequence-based control. We visualized the resulting dendrograms using the scipy^66^ package.

#### Goodness of clustering metrics

We selected three standard clustering metrics to evaluate the quality of inferred cluster identities compared to the true family classification given by the expert-annotated labels. The metrics we selected were required to be bounded between 0 and 1, to be invariant to the number of families and clusters present in the data, and to be symmetric. We therefore selected Adjusted Mutual Information, Adjusted Rand Index, and Fowlkes Mallows Score for these properties. The definition, usage, and properties of these metrics are described elsewhere^67^.

### Fine-tuning to generalize GFP function prediction for protein engineering

#### Collecting fluorescent protein homologs with EBI JackHMMer

We sourced Fluorescent Protein sequences from the literature and public databases (Interpro IPR011584, IPR009017; PFAM:PF01353,PF07474 and^68, 69^). We were left with 1135 sequences total after cleaning sequences longer than 1000 AAs or containing invalid letters. We the used the python-Levenshtein package to compute distances from sfGFP. Looking to source sequences of varying dissimilarities from sfGFP, we sampled <100 sequence proximity-ordered subsets, first selecting the most distant subset with probability proportional to the distance from sfGFP, and then continuing iteratively until the most similar set was apportioned (this subset contained less than 100 sequences unlike the rest). We then used the EBI JackHMMer web server^70^ to batch search each subset, with no more than 20 iterations. Search was stopped after more than 100,000 sequences were discovered or the search converged. We expected this batched approach to generate unique hits within each subset, but we found that after cleaning to remove long (>500 AAs) or invalid seqs, and dropping duplicates with preference to the subset nearest sfGFP, almost all the sequences were discovered by the most nearby subset search. We continued with these 32,225 sequences.

With the goal of establishing a validation set which was measuring something closer to extrapolation, we took these sequences and recomputed distance with sfGFP. We selected a distance-biased 10% “out of domain” validation set by sampling with probability proportional to the 4th power of the distance (strongly weighting distant examples). We also selected a 10% “in-domain” validation set uniformly randomly from the remaining sequences. This left 25,781 training sequences.

#### Model finetuning with extant GFP sequences

We loaded the weights learned by UniRep in the exact same architecture as before, but replacing the final layer, which previously predicted the next character, with a randomly initialized feed-forward layer with a single output and no non-linearity. As a control, we initialized the same randomly. We trained both models with exactly the same procedure: low learning rate (.00001), 128 batch size, and only partially feeding forward along the sequence, stopping prediction after 280 amino acids, with full back propagation rather than truncated as during the UniRef50 training process. This was determined by the computational constraint to fit the unrolled recurrent computational graph into GPU memory; however, we expected this was sufficiently long to capture the context of GFP because the vast majority of well-characterized fluorescent proteins are shorter than 280 AA. Both models trained for ∼13,000 weight updates, corresponding to ∼65 epochs, and ∼1.5 days wall clock time on 1 Nvidia Volta GPU. Stopping was determined by computational constraints, not overfitting as measured by increasing validation set loss, which never occurred.

#### Training LASSO regression on representation featurized sequences from the fitness landscape of avGFP

Sequences from Sarkisyan et. al (2016)^37^ were featurized using a given representation (e.g. UniRep). A sparse LASSO linear regression model was trained using this representation as input. A range of 20 L1 penalties, log-spaced over 6 orders of magnitude, was scanned using 10-fold cross validation. The selected level of regularization was set to be the strongest (most regularizing) penalty that had statistically equal out-of-sample error to the penalty with lowest out-of-sample error across the 10 folds.

#### Baseline selection for retrospective fluorescent protein sequence discovery task

Doc2Vec is a standard tool for numerically representing text documents in natural language processing. Yang *et al.* (2018) use this simple approach to represent protein sequences^21^ and achieve good supervised function prediction performance with simple models based on their representation. We also found these Doc2Vec representations to be especially appropriate baselines as they were originally validated, in part, by reaching state-of-the-art performance on a rhodopsin absorption wavelength prediction task, which bears similarity to fluorescent prediction task at hand.

We did not consider Rosetta an appropriate baseline here for three reasons: 1) Rosetta provides measures of stability, which does not completely define function, 2) the computational requirements to mutate and relax a reference structure for >32,000 sequences, let alone de novo fold, are impractical, and 3) in general, we would not expect to have structures for sequences we have yet to discover, meaning we would need to rely on the distant avGFP as a template.

We additionally could not apply simple linear regression or Naive Bayes approaches here as those methods require fixed length inputs whereas the protein length sequences examined here were variable length.

#### Processing well-characterized GFP sequences from FPBase

A raw collection of 452 fluorescent proteins was sourced from FPbase.org ^39^ (Collection “FP database” available at https://www.fpbase.org/collection/13/ as of Nov 2, 2018). This was then filtered as follows:

1. Sequences shorter than 200 amino acids or longer than 280 amino acids were removed.
2. Remaining sequences were then multiply aligned using ClustalW (gap open penalty=10, gap extension penalty=0.1) ^71^. BLOSUM62 was used as the scoring matrix.
3. Using these aligned sequences, we computed their pairwise minimum edit distance (insertions/deletions counting as one edit). Sequences that were more than 50 mutations away from their nearest neighbor were removed. These were enriched for circularly permuted FPs and tandem FPs.
4. Remaining that had an average edit distance >180 mutations (length of avGFP = 238) to all other sequences were also removed.
5. Where possible, His tags and likely linker sequences were manually removed, and start Methionine amino acids were added to all sequences that did not have one.
6. At this point, the major phylogenetic clusters of Anthozoan (corals/anemones) and Hydrozoan (Jellyfish) FPs remained including engineered variants. Note that hydrozoan and anthozoans are almost entirely different, often sharing as little as 30% sequence similarity. The hydrozoan clade was almost entirely composed of FPs found in *Aequorea victoria* or engineered versions thereof. The set of sequences for the retrospective sequence discovery analysis was therefore set to be all natural or engineered green (emission wavelength between 492 nm and 577 nm) fluorescent proteins of *Aequorean* descent.
7. This set of sequences included well known engineered variants of avGFP including EGFP, mVenus, superfolder GFP, mCitrine, and Clover.

### Exploratory analysis and data visualization

#### PCA of amino acid embeddings

We extracted embedding vectors for each amino acid from the trained UniRep model. We performed PCA as implemented in the sklearn library and used the three first principal components for the visualization.

#### t-SNE of organism proteomes

We obtained 53 reference proteomes (Supp. Table 1) of model organisms from UniProt Reference Proteomes. We used the UniRep trained model to obtain representations for each of the proteins in each proteome. We then averaged all the proteins in each proteome to obtain the representation for the “average protein” for each of the organisms. We used t-SNE^24^ - a common technique for visualizing high-dimensional datasets (as implemented in the sklearn library, with perplexity=12) to obtain a 2D projection.

#### PCA of conserved protein organism relationships

To allow embedding of new points into the projected space, we computed a PCA using a subset of better characterized reference proteomes (as labelled in Supp. Fig. S2). We manually sourced 5 conserved sets of proteins from Humans, S. cerevisiae, and D. rerio using the OrthoDB ^72^ as well as manual inspection and UniProt characterization data. These proteins were: dihydrofolate reductase (DHFR), methylenetetrahydrofolate reductase (MTHFR), nucleotide excision repair protein (ERCC2), elongation factor (EFTU), Heat-Shock protein 70 (HSP701a). We projected the representations for each of these variants into the space given by the first two Principal Components from the model organism PCA and drew the vector from S. cerevisiae to Human, which we translated so the base sits on the S. cerevisiae variant for each protein.

#### Correlation of representation with structural features

We computed the full sequence of hidden states at every position of 3720 single-domain proteins from the SCOP database ^29^ (for which we had secondary structure annotation for each of the positions obtained from PDB ^73^). We proceeded to calculate correlations between elements of these hidden sequences (neurons) and secondary structure. Neuron 341 was a strong helix *and* beta-sheet detector, obtaining 0.33 correlation with alpha-helicity and −0.35 correlation with beta-sheets.

We then cut out all continuous helices and beta-sheets with surrounding context (including [30% * the length of the helix/sheet] context amino acids on each side) from the proteins in the database to look at the average activation of the neuron at each relative position in the helix/sheet, obtaining ∼14000 alpha helices and 20,000 beta-sheets.

We then similarly computed all hidden states for 1,500 randomly sampled proteins with available structures from the Protein Data Bank (PDB). We attempted to compute DSSP ^74^ annotations for each structure, keeping only those sequences and corresponding structures such that the DSSP calculation executed without error and DSSP secondary structure amino acid indices lined up with the primary amino acid indices. These calculations succeeded for 448/1,500 structures. After exploratory correlational analysis of various structural features, we decided to focus on solvent accessibility. Without explicit regularization on the hidden state of the LSTM, we felt it was likely that the representation was entangled, with multiple neurons possibly encoding the same biophysically relevant features. We therefore also learned, in a supervised manner, simple and sparse linear combination of neurons that were predictive of solvent accessibility. To train this model we used a position by hidden-sequence dimension matrix as the feature matrix, and solvent accessibility as a response. These were both input to LASSO and the strength of L1 regularization was selected using 10 fold cross validation.

Single neuron activations or linear combinations thereof were visualized on protein structures using the NGLview Python package^75^.

## List of Supplementary materials

**Supplemental Figure 1.** Growth in sequence databases.

**Supplemental Table 1.** Reference proteomes used in the organism analysis in Fig. 2b.

**Supplemental Figure 2.** Single-protein vector arithmetic in UniRep representation space.

**Supplemental Table 2.** Unirep achieves competitive results on homology detection as measured by ROC-AUC and ROC50-AUC (sorted by ROC score).

**Supplemental Figure 3.** A representative clustering of Cytochrome-oxidase family enzymes from HOMSTRAD (Fig. 2d) further illustrates this result by reconstructing the correct monophyletic grouping of the true labels, where a sequence distance-based clustering fails.

**Supplemental Figure 4.** OXBench unsupervised homology detection task, all results.

**Supplemental Figure 5.** HOMSTRAD unsupervised homology detection task, all representations results.

**Supplemental Figure 6**. Learned from a collection of PDB secondary structures, a linear combination of hidden state neurons identifies solvent accessible regions in the structure of Bovine Rhodopsin GPCR (PDB:1F88, left) oriented with the extracellular domain upwards (Methods).

**Supplemental Figure 7.** Baseline representations.

**Supplemental Table 3.** Representation fusions (concatenations) analyzed.

**Supplemental Table 4.** Regression results - test set metrics.

**Supplemental Table 5.** Regression results - validation set metrics.

**Supplemental Figure 8.** Validation scores for main text Figure 3e, 17 DMS protein stability prediction datasets.

**Supplemental Figure 9.** Linear model on top of UniRep is simpler (has fewer parameters) than the the same with basic One-Hot-Encoding standardly used for sequences if the sequence is longer than 95aa.

**Supplemental Figure 10.** Variant Effect and stability generalization tasks; hypothesized mechanism for transfer performance.

**Supplemental Figure 11.** Efficiency curves for all baseline representations: recall (a) and maximum brightness (b).

**Supplemental Figure 12.** UniRep cost savings in GFP protein engineering tasks.

**Supplemental Table 6.** Analysis datasets and tasks.

**Supplemental Figure 13.** UniRep is a generative model of protein sequences.

**Supplemental Table 7.** Data augmentation of UniRep. Supplemental Table 8. All the models we evaluated.

**Supplemental Data 1.** All test set results in .xlsx format: https://s3.amazonaws.com/unirep-public/supp_data_1_test_set_full.xlsx

**Supplemental Data 2**. All validation set results in .xlsx format: https://s3.amazonaws.com/unirep-public/supp_data_2_val_set_full.xlsx

**Supplemental Data 3.** All test set results graphically presented in .pdf format: https://s3.amazonaws.com/unirep-public/supp_data_3_graph_test_set.pdf

## Supplementary Materials

**Supplemental Figure 1.**
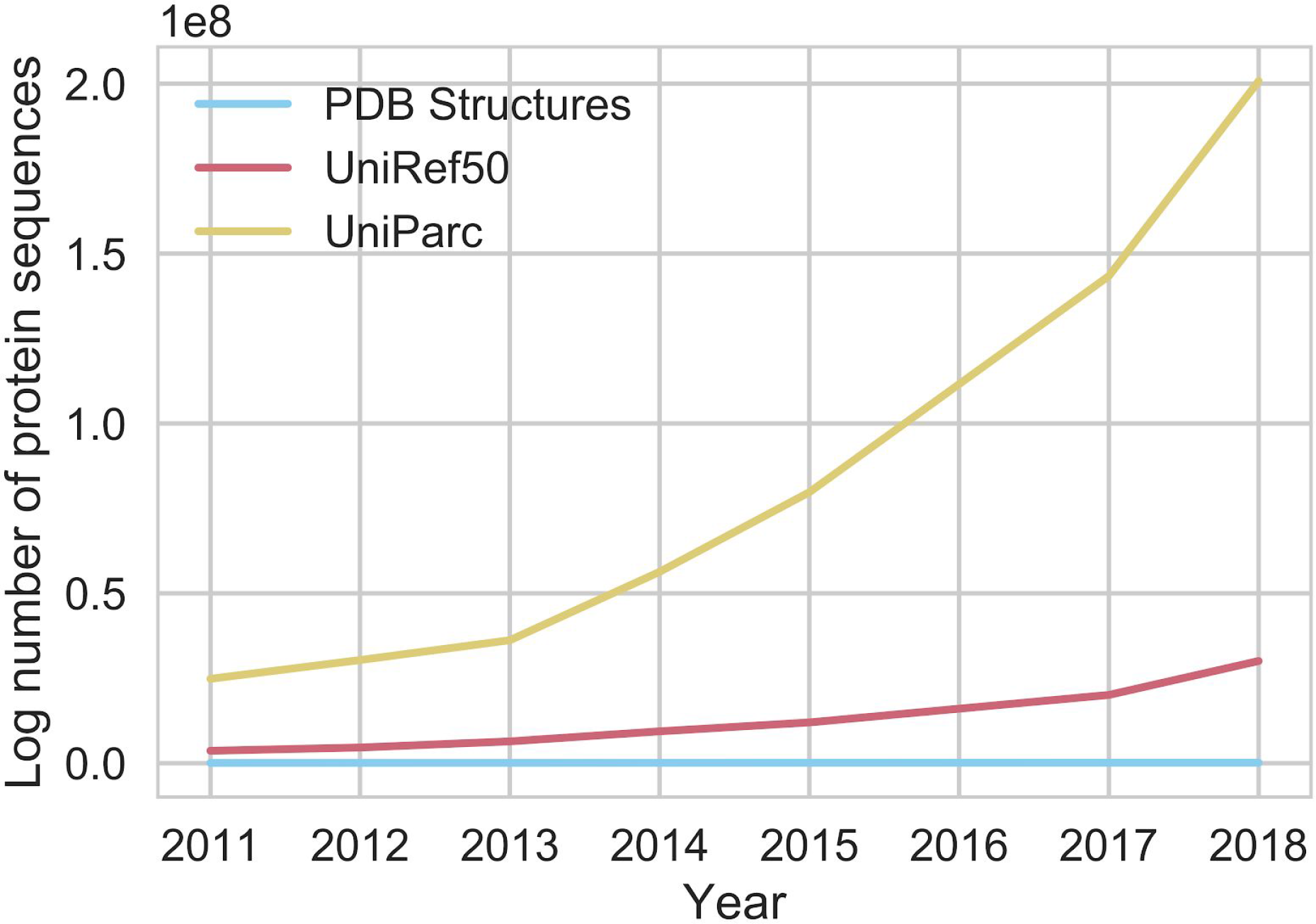
Growth in sequence databases.

**Supplemental Table 1.** Reference proteomes used in the organism analysis in Fig. 2b. Species name and common/ subspecies/ strain in parentheses.

| Organism Short Name | Domain | Specific Name |
| --- | --- | --- |
| *Halobacterium salinarum* | Archaea | Archaea |
| *Haloferax volcanii* | Archaea | Archaea |
| *Methanococcus maripaludis* | Archaea | Archaea |
| *Methanosarcina acetivorans* | Archaea | Archaea |
| *Sulfolobus solfataricus* | Archaea | Archaea |
| *Thermococcus kodakarensis* | Archaea | Archaea |
| *Escherichia coli* (K12) | Bacteria | Bacteria |
| *Aliivibrio fischeri* | Bacteria | Bacteria |
| *Azotobacter vinelandii* | Bacteria | Bacteria |
| *Bacillus subtilis* (168) | Bacteria | Bacteria |
| *Cyanothece* (PCC7822) | Bacteria | Cyanobacteria |
| *Mycoplasma genitalium* | Bacteria | Bacteria |
| *Mycobacterium tuberculosis* | Bacteria | Bacteria |
| *Prochlorococcus marinus* | Bacteria | Cyanobacteria |
| *Streptomyces coelicolor* (A32) | Bacteria | Bacteria |
| *Synechocystis* (PCC 6803 Kazusa) | Bacteria | Cyanobacteria |
| *Caenorhabditis elegans* | Eukarya | Animalia |
| *Drosophila melanogaster* (fruit fly) | Eukarya | Animalia |
| *Homo sapiens* (human) | Eukarya | Mammalia |
| *Mus musculus* (mouse) | Eukarya | Mammalia |
| *Saccharomyces cerevisiae* (yeast) | Eukarya | Fungi |
| *Anolis carolinensis* | Eukarya | Animalia |
| *Aspergillus nidulans* | Eukarya | Fungi |
| *Arabidopsis thaliana* | Eukarya | Plantae |
| *Cavia porcellus* (guinea pig) | Eukarya | Mammalia |
| *Gallus gallus domesticus* (chicken) | Eukarya | Animalia |
| *Coprinopsis cinerea* | Eukarya | Fungi |
| *Bos taurus* (cow) | Eukarya | Mammalia |
| *Chlamydomonas reinhardtii* | Eukarya | Eukarya |
| *Cryptococcus neoformans* | Eukarya | Fungi |
| *Canis familiaris* (dog) | Eukarya | Mammalia |
| *Emiliana huxleyi* | Eukarya | Eukarya |
| *Macaca mulatta* (macaque) | Eukarya | Mammalia |
| *Zea mays* (corn) | Eukarya | Plantae |
| *Heterocephalus glaber* (naked mole rat) | Eukarya | Mammalia |
| *Neurospora crassa* | Eukarya | Fungi |
| *Oryzias latipes* | Eukarya | Animalia |
| *Physcomitrella patens* | Eukarya | Plantae |
| *Columba liva* (pigeon) | Eukarya | Animalia |
| *Sus scrofa* (pig) | Eukarya | Mammalia |
| *Pristionchus pacificus* | Eukarya | Animalia |
| *Oryza sativa* (Rice strain japonica) | Eukarya | Plantae |
| *Schizosaccharomyces pombe* | Eukarya | Fungi |
| *Tetrahymena thermophila* | Eukarya | Eukarya |
| *Thalassiosira pseudonana* | Eukarya | Eukarya |
| Ustilago maydis (corn smut) | Eukarya | Fungi |
| *Xenopus tropicalis* | Eukarya | Animalia |
| *Danio rerio* | Eukarya | Animalia |
| Phage lambda | Virus | Virus |
| SV40 | Virus | Virus |
| T4 phage | Virus | Virus |
| T7 phage | Virus | Virus |
| Vaccinia virus (Copenhagen) | Virus | Virus |

**Supplemental Figure 2.**
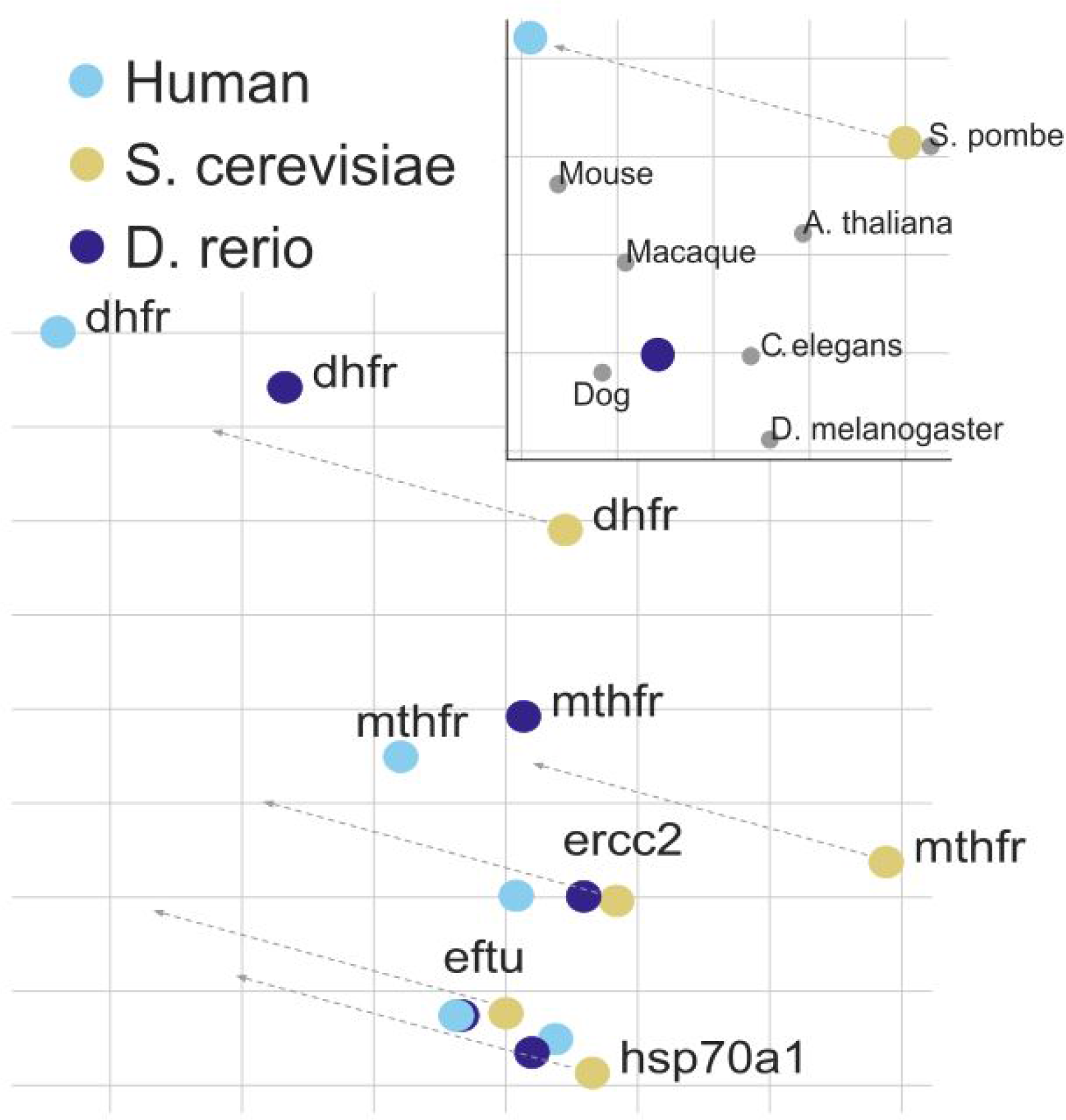
Single-protein vector arithmetic in UniRep representation space. We suspected our organism vector clustering success may be explained by learning a measure of proteome content (e.g. abundance of various protein types). Surprisingly, after sourcing 5 proteins conserved across 3 model organisms, we identify a common direction of variance, from Baker’s Yeast to Human in the PCA projection space, which corresponds to the vector from Yeast to Human proteome representations, suggesting that organisms may have an arithmetic relationship in the representation space similar to that observed in Word Vectors ^76^. Note that the direction of the vectors is invariant from the PCA in the upper right to the bottom left, but the length of the vector is meaningless in the bottom left. PC1 is the x-axis of both plots, PC2 is the y axis of both plots.

**Supplemental Table 2.** Unirep achieves competitive results on homology detection as measured by ROC-AUC and ROC50-AUC (sorted by ROC score). UniRep with RandomForest top model with Bayesian Hyperparameter optimization (Methods) achieves competitive performance with published sequence-only remote homology detection methods (Håndstad, 2007) on two most frequently used benchmark datasets.

| SCOP 1.67 Superfamily Level<br>Remote Homology Detection |  |  | SCOP 1.67 Fold Level Remote<br>Homology Detection |  |  |
| --- | --- | --- | --- | --- | --- |
|  | ROC | ROC50 |  | ROC | ROC50 |
| LA-Kernel | 0.92 | 0.69 | GPKernel | 0.84 | 0.51 |
| GPKernel | 0.9 | 0.59 | LA-Kernel | 0.83 | 0.5 |
| UniRep | 0.89 | 0.49 | Mismatch | 0.81 | 0.47 |
| Mismatch | 0.88 | 0.54 | UniRep | 0.81 | 0.44 |
| GPextended | 0.87 | 0.54 | GPextended | 0.75 | 0.37 |
| eMOTIF | 0.86 | 0.55 | SVM-Pairwise | 0.72 | 0.36 |
| SVM-Pairwise | 0.85 | 0.56 | eMOTIF | 0.7 | 0.31 |
| GPBoost | 0.8 | 0.38 | GPBoost | 0.69 | 0.3 |
| PSI-BLAST | 0.57 | 0.17 | PSI-BLAST | 0.5 | 0.01 |

**Supplemental Figure 3.**
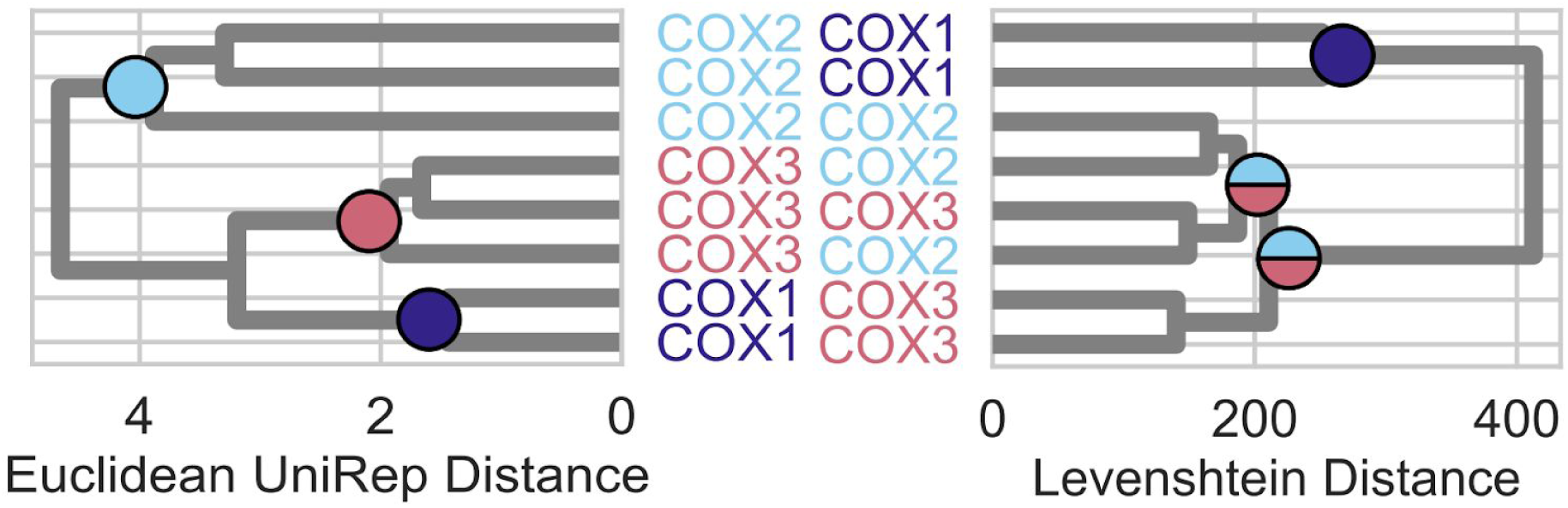
Euclidean distance in UniRep space resolves protein clusters where generalized minimum edit distance fails. A representative clustering of Cytochrome-oxidase family enzymes (COX) 1, 2, and 3 from HOMSTRAD (Fig. 3e) further illustrates this result by reconstructing the correct monophyletic grouping of the true labels, where a sequence distance-based clustering fails.

**Supplemental Figure 4.**
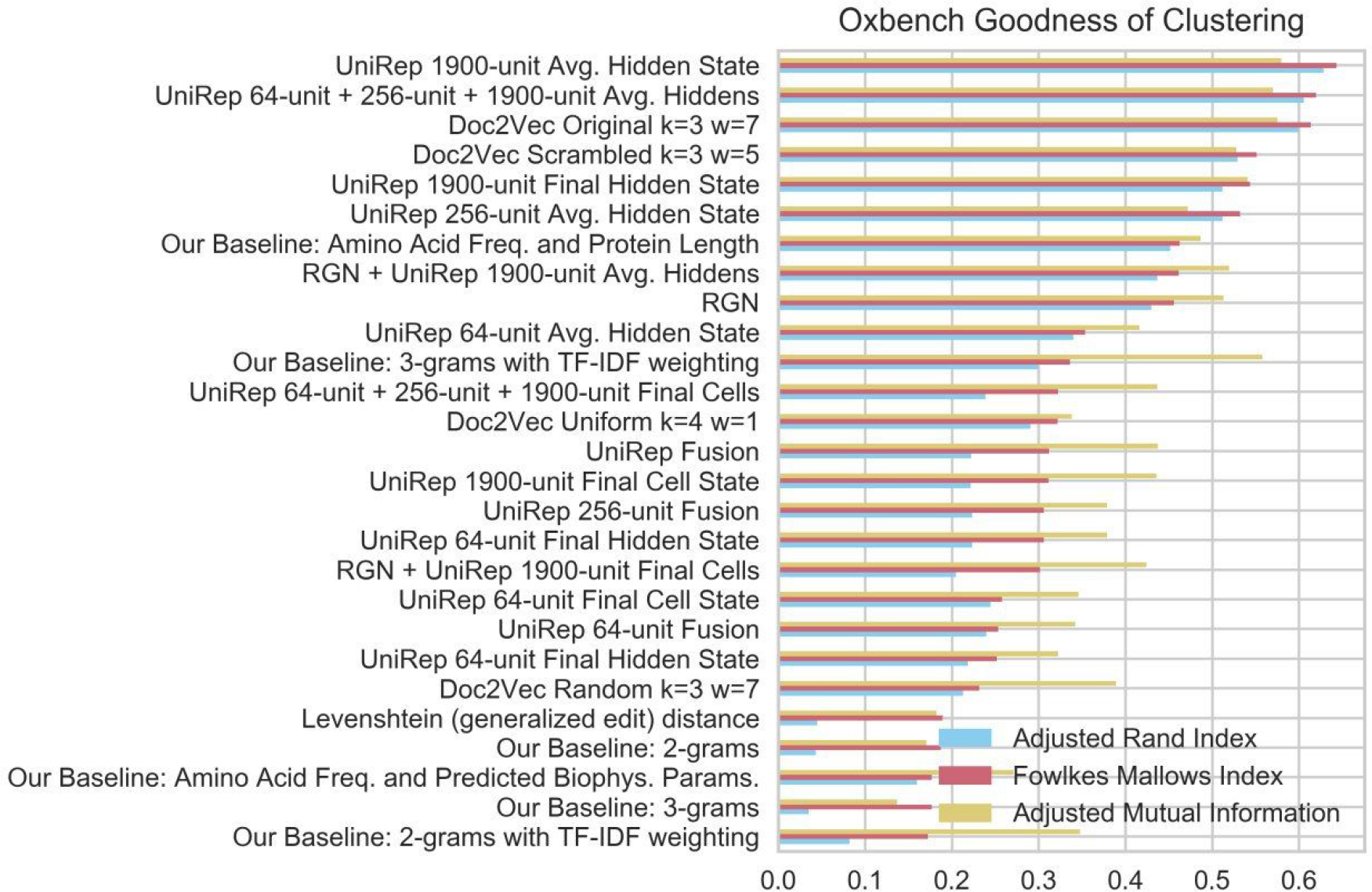
OXBench unsupervised homology detection task, all results.

**Supplemental Figure 5.**
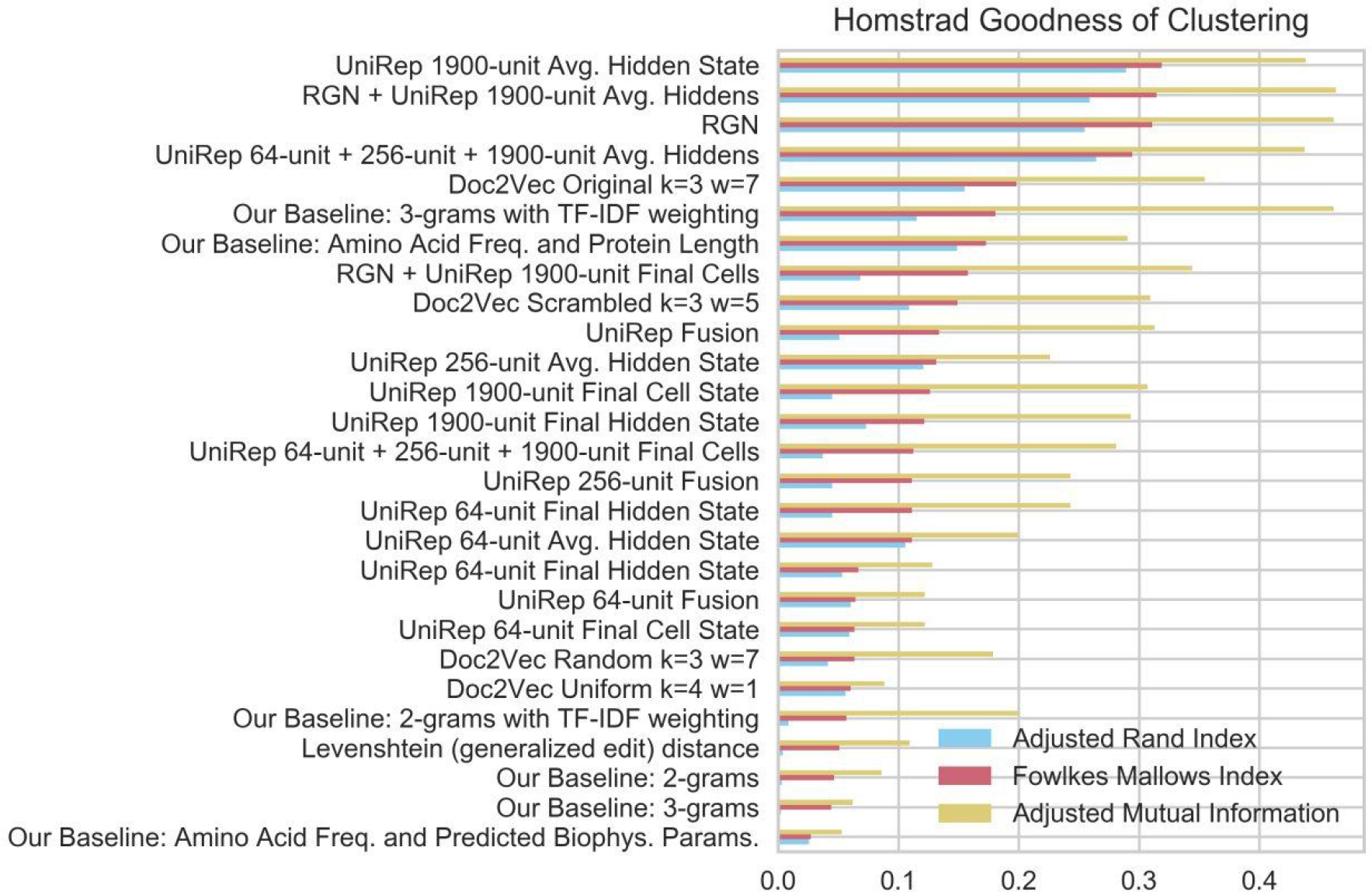
HOMSTRAD unsupervised homology detection task, all representations results.

**Supplemental Figure 6.**
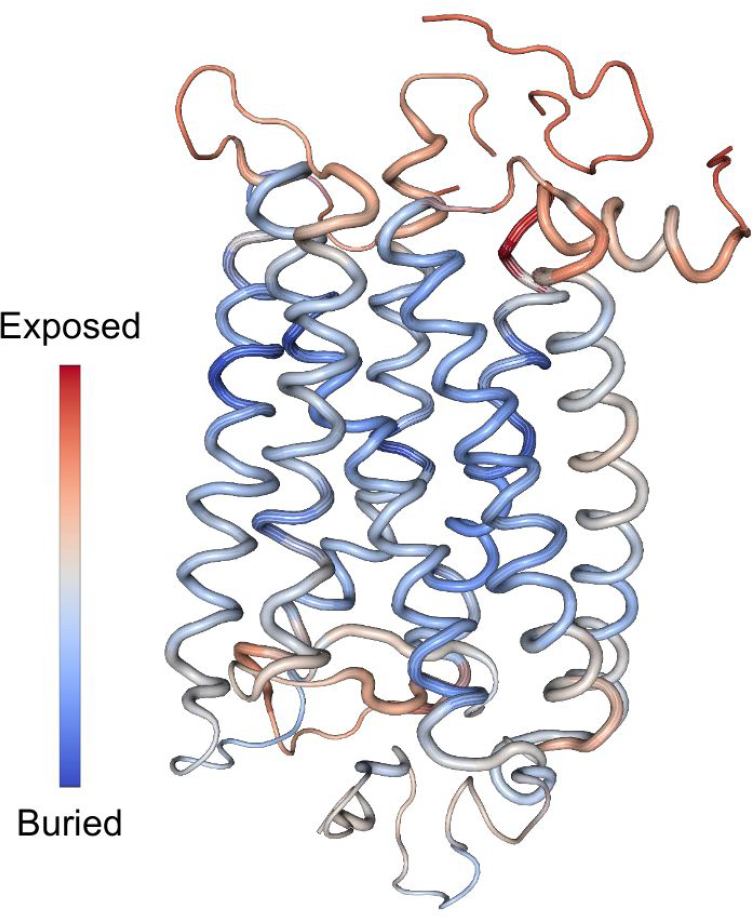
Learned from a collection of PDB secondary structures, a linear combination of hidden state neurons identifies solvent accessible regions in the structure of Bovine Rhodopsin GPCR (PDB:1F88, left) oriented with the extracellular domain upwards (Methods). Can predict solvent accessibility with a Pearson correlation of 0.38.

**Supplemental Figure 7.**
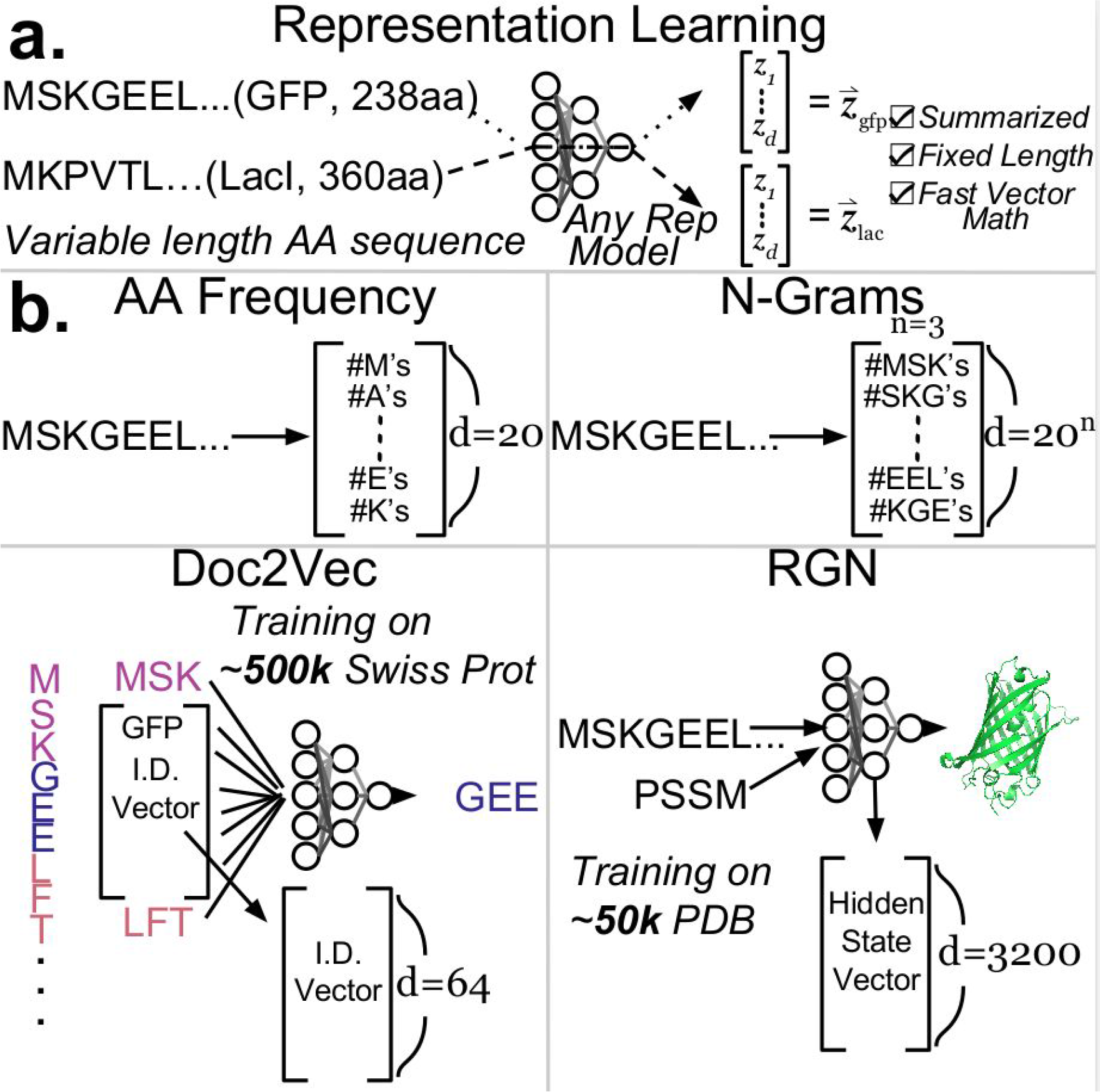
Baseline representations. Representation learning inputs primary amino acid sequences and outputs fixed length vector representations. **a.** Schematic of representation learning in general. **b.** 4 baseline methods illustrated: simply counting amino acid occurrences (upper left) and occurrences of k-mers (upper right), Doc2Vec embeddings learned by Feed-Forward prediction of a central k-mer given the external context k-mers (lower left) ^21^, Recurrent Geometric Network hidden state, learned by recurrent processing of input sequences to predict crystal structure (lower right) ^16^.

**Supplemental Table 3.**
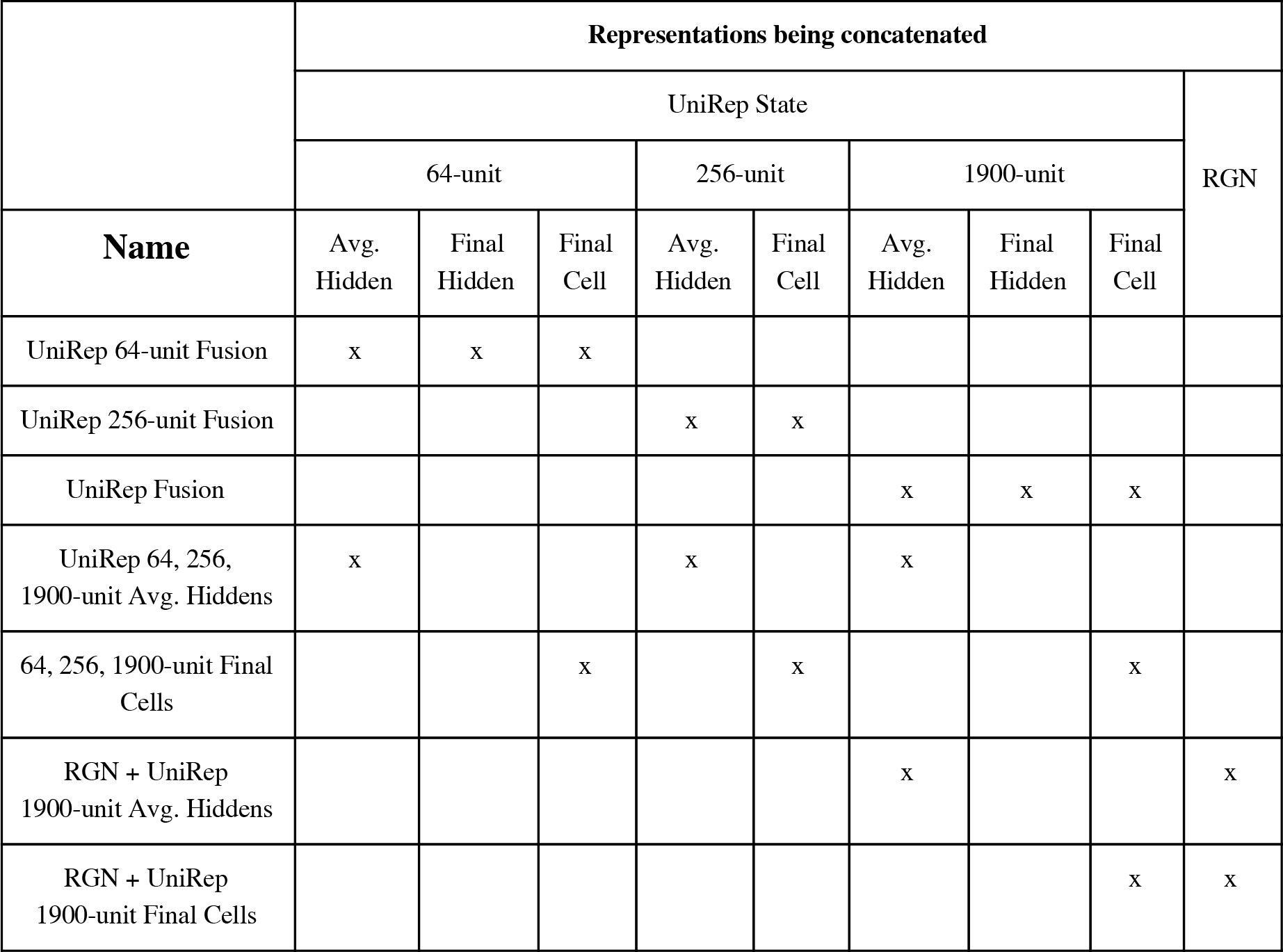
Representation fusions (concatenations) analyzed.

|  | Representations being concatenated |  |  |  |  |  |  |  | RGN |
| --- | --- | --- | --- | --- | --- | --- | --- | --- | --- |
|  | UniRep State |  |  |  |  |  |  |  |  |
|  | 64-unit |  |  | 256-unit |  | 1900-unit |  |  |  |
| Name | Avg.<br>Hidden | Final<br>Hidden | Final<br>Cell | Avg.<br>Hidden | Final<br>Cell | Avg.<br>Hidden | Final<br>Hidden | Final<br>Cell |  |
| UniRep 64-unit Fusion | x | x | x |  |  |  |  |  |  |
| UniRep 256-unit Fusion |  |  |  | x | x |  |  |  |  |
| UniRep Fusion |  |  |  |  |  | x | x | x |  |
| UniRep 64, 256,<br>1900-unit Avg. Hiddens | x |  |  | x |  | x |  |  |  |
| 64, 256, 1900-unit Final<br>Cells |  |  | x |  | x |  |  | x |  |
| RGN + UniRep<br>1900-unit Avg. Hiddens |  |  |  |  |  | x |  |  | x |
| RGN + UniRep<br>1900-unit Final Cells |  |  |  |  |  |  |  | x | x |

**Supplemental Table 4.** Regression results - test set metrics, with lowest MSE model or model class compared to the 2nd lowest MSE model or model class *p < 0.05, **p < 0.01, ***p<0.001 (Welch’s t-test for significance), standard deviations obtained through 30x 50% validation/test set resampling. Validation set metrics can be found in Supp. Table 5. This table includes an extension of our analysis to 4 small datasets compiled previously for protein phenotype prediction using Doc2Vec representations (first 4 columns) ^21^. We observed widely variable results and statistically insignificant results (caused by underpowered validation and test set), with UniRep or one of the baselines we developed outperforming previous state-of-the-art ^21^ (here and in Supp. Table S5), which underscored the importance of adequate data size for accurate estimation of performance.

|  |  | Cytochrome<br>P450<br>Thermostability | Rhodopsin<br>Peak<br>Absorption<br>Wavelength | Epoxide<br>Hydrolase<br>Enantio<br>selectivity | Channel<br>rhodopsin<br>Membrane<br>Localization | TEM-1 Beta-<br>lactamase | Ubiquitin<br>(E1<br>Activity) | Protein G<br>(IgG<br>domain) |  |
| --- | --- | --- | --- | --- | --- | --- | --- | --- | --- |
| UniRep Fusion |  | **15.8** | **499** | 189 | 1.25 | **0.0545\*\*\*** | **0.0421\*\*\*** | **0.0233\*\*\*** |  |
| Our Best Baseline |  | 21.7 | 571 | **93.2** | **0.912** | 0.074 | 0.052 | 0.054 |  |
| RGN |  | 24.4 | >2000 | >1000 | 1.61 | 0.0904 | 0.054 | 0.0977 |  |
| Best Doc2Vec |  | 18.1 | 530 | 95.7 | 1 | 0.0881 | 0.0625 | 0.0724 |  |
|  |  |  |  |  |  |  |  |  | Spearman<br>Rank |
|  | HSP90 | Amino<br>glycosidase<br>(Kka2) | Pab1<br>(RRM<br>domain) | PSD95<br>(Pd3<br>domain) | Ubiquitin | Yap65<br>(WW<br>domain) | Stability: 17<br>DMS datasets<br>combined | Stability: *De*<br>*Novo* Designed<br>mini proteins | Correlation<br>Stability: *De*<br>*Novo* Design<br>Rounds |
| UniRep Fusion | **0.0258\*\*\*** | **0.11\*\*** | **0.0265\*\*\*** | **0.0208\*\*\*** | **0.0323\*\*\*** | **0.0415\*\*\*** | **0.0304\*\*\*** | **0.179\*\*\*** | **$\rho=0.59***$** |
| Our Best Baseline | 0.0344 | 0.115 | 0.0435 | 0.041 | 0.0515 | 0.0662 | 0.0398 | 0.201 | - |
| RGN | 0.0579 | 0.14 | 0.0596 | 0.0438 | 0.0601 | 0.0639 | 0.0338 | 0.189 | - |
| Best Doc2Vec | 0.0579 | 0.132 | 0.0495 | 0.046 | 0.064 | 0.0772 | 0.0473 | 0.258 | - |
| | | | | | | | | Rosetta Total<br>Energy | $\rho=0.42$ |

**Supplemental Table 5.**
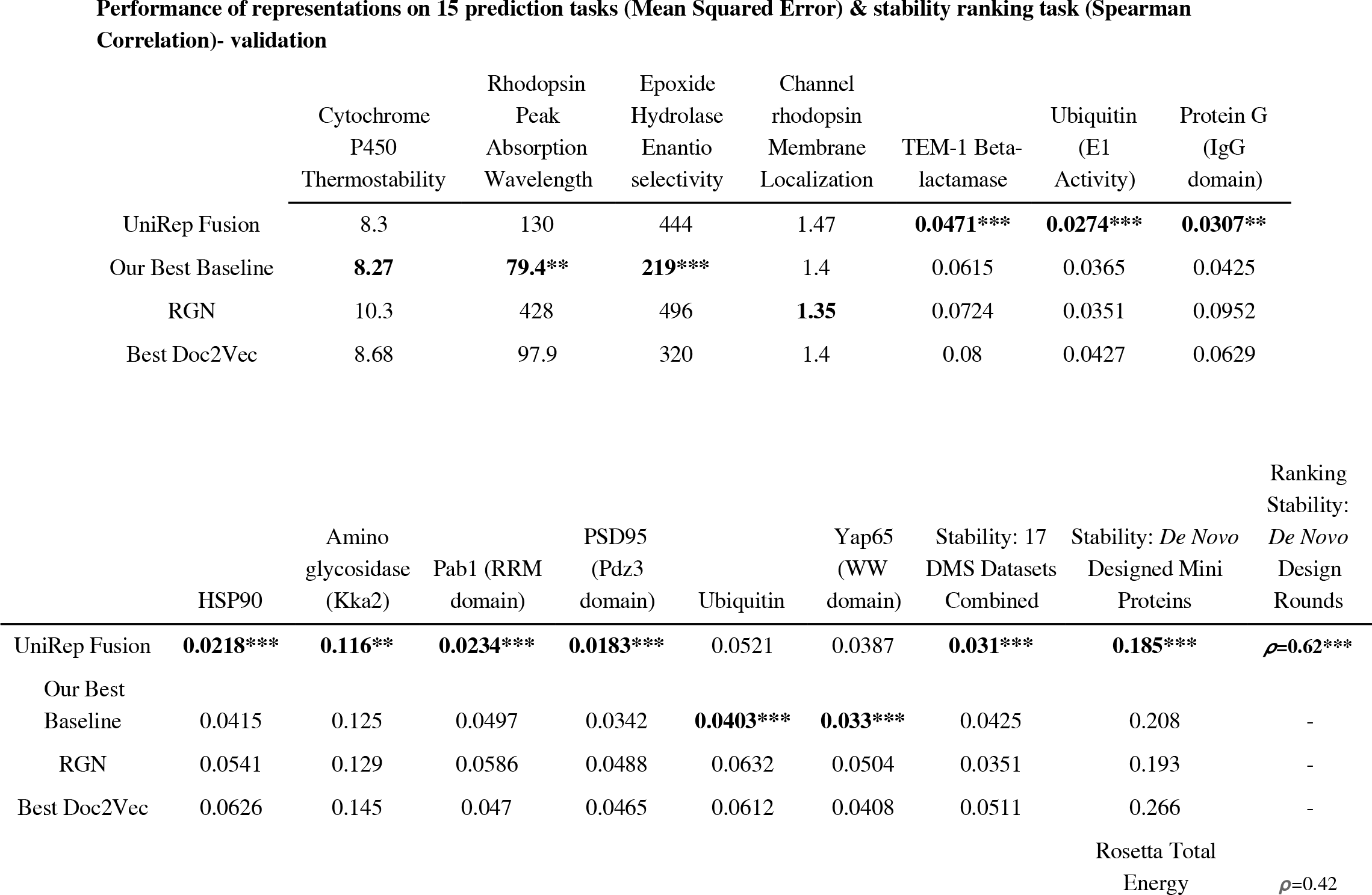
Regression results - validation set metrics, with lowest MSE model or model class compared to the 2nd lowest MSE model or model class *p < 0.05, **p < 0.01, ***p<0.001 (Welch’s t-test for significance), standard deviations obtained through 30x 50% validation/test set resampling. Test set metrics and explanation of the first 4 column tasks can be found in Supp. Table 4.

**Supplemental Figure 8.**
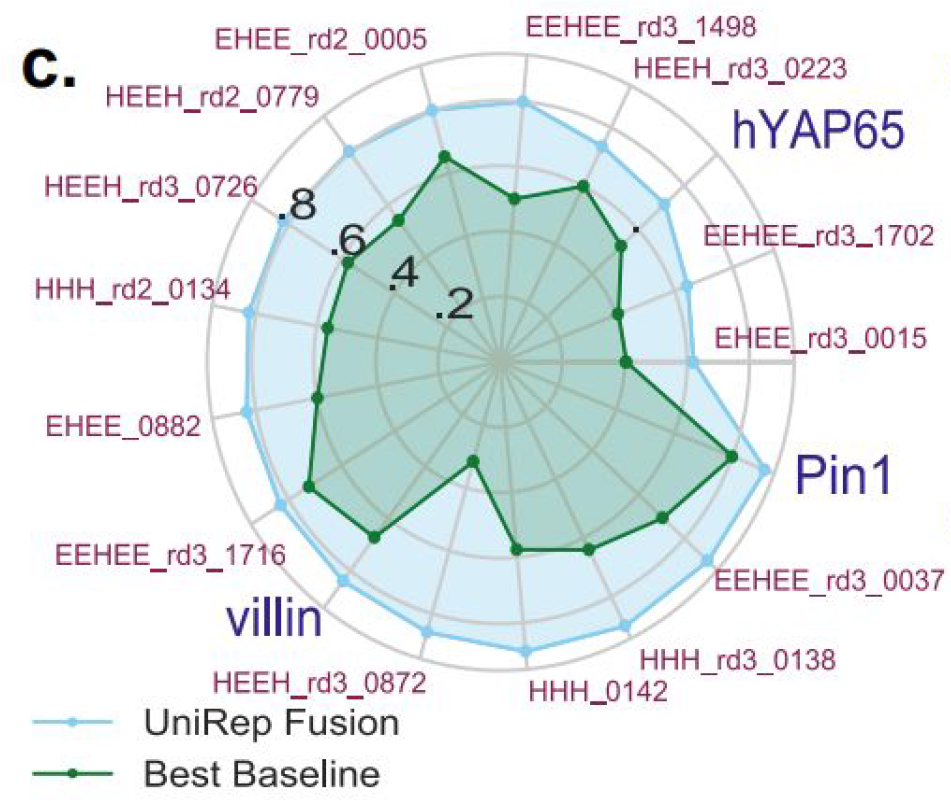
Validation scores for main text Figure 3e, 17 DMS protein stability prediction datasets.

**Supplemental Figure 9.**
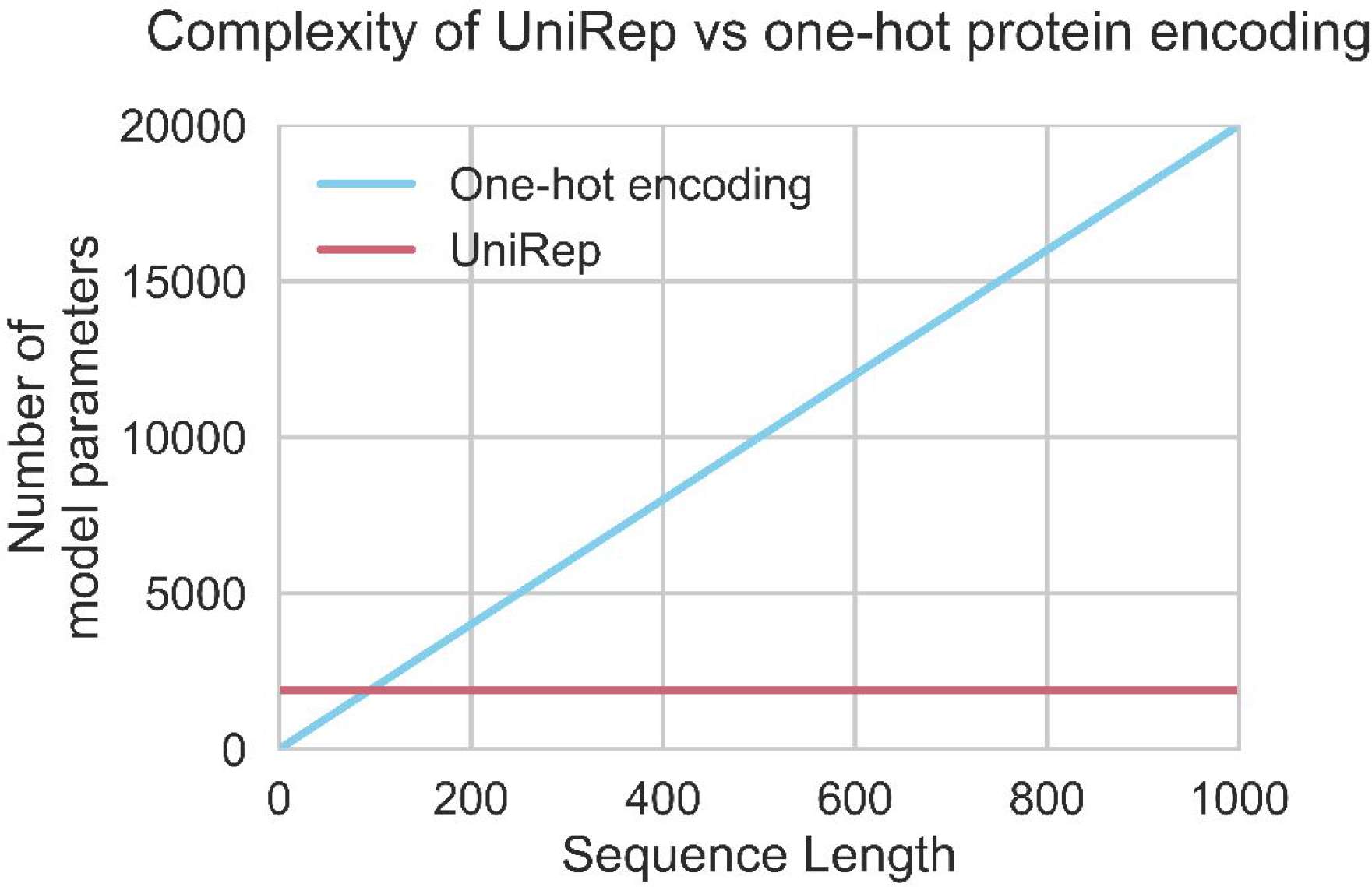
Linear model on top of UniRep is simpler (has fewer parameters) than the same model using a standard One-Hot-Encoding if the sequence is longer than 95aa.

**Supplemental Figure 10.**
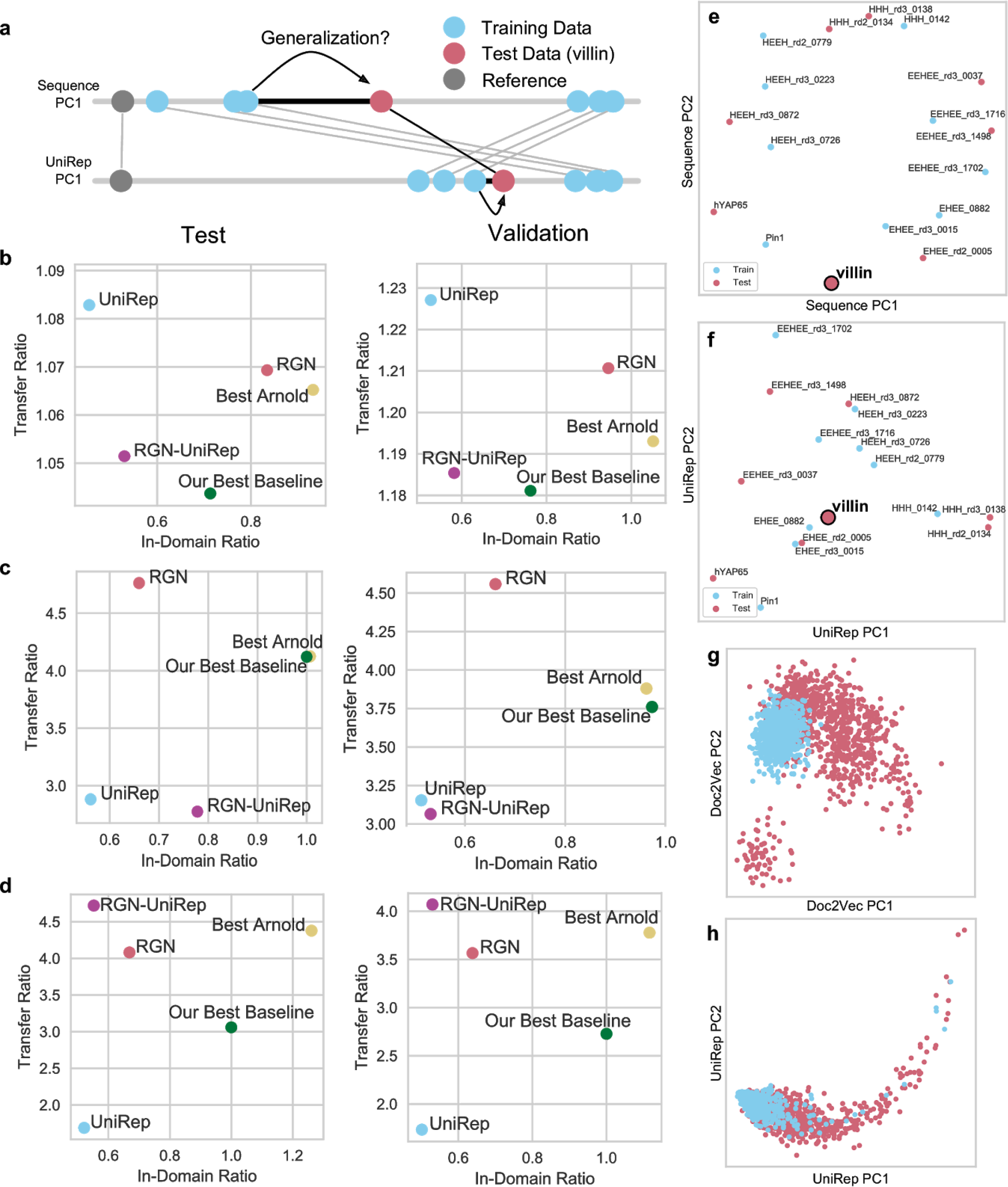
Variant Effect and stability generalization tasks; hypothesized mechanism for transfer performance. We used a well-established generalization scoring methodology from work in machine learning^61^. It quantifies the error of local, “In-Domain” predictions as well as generalization error, which they call “Transfer”, relative to a baseline (Methods); *lower is better*. Unless stated otherwise, we use the Leave One Protein Out (LOPO) procedure, withholding one protein from the set at a time, training a model on all but that one protein, evaluating it on the single withheld protein, and taking an average of these generalization errors (Methods) ^35^. **a.** Generalization is making accurate predictions on sequences that are distant from the training data. Here, the training data (blue), a distant reference sequence (grey) and the the test data (villin) are shown on the first Principle Coordinate of an MDS of the Levenshtein sequence distance. In sequence space, villin is far from the nearest training point, which makes generalization challenging. In UniRep space, shown here as the first Principal Component, the training and test data are rearranged so villin is much closer to the nearest training point, thereby enabling generalization. **b.** Generalization performance on the LOPO generalized variant effect prediction task as measured by In-Domain vs Transfer ratios for test set (left) and validation set (right). We used the function prediction dataset from Figure 3 with 8 proteins with 9 distinct functions measured in separate experiments ^35, 77^. It was previously shown that some of the regions most vulnerable to deleterious single mutations are functional and highly conserved or co-conserved in evolution ^78^. Standard approaches therefore rely strongly on co-evolutionary data ^15, 35, 77^ or even structural data ^35^, which implicitly demarcate functional residues. Because it has neither of these as inputs and was trained on a corpus with at most 50% similarity between sequences, UniRep should find it challenging to identify such residues. Nevertheless, UniRep performs best in-Domain, successfully identifying functionally-important positions from some labeled mutation data for a given protein. However, as expected, UniRep does not generalize well to proteins for which it had no labeled training data, at least in this case. When we tested fusing UniRep to the RGN (RGN-Fusion), which was trained on a form of evolutionary data (PSSMs) and predicts protein structure, we see a boost in performance suggesting a good trade-off between In-Domain and Transfer Error. **c.** Generalization performance on the LOPO generalized DMS stability task, measured by In-Domain and Transfer ratios for test set (left) and validation set (right). Unlike variant effect, stability is a property consistent to all proteins. There were 3 natural and 14 *de novo* designed wildtypes. Using the same LOPO procedure as described above, we found that UniRep outperformed all baselines at generalization, and that the RGN no longer offered meaningfully complementary information, RGN-Fusion performing approximately as well as UniRep. This suggests UniRep does enable generalization of universal protein characteristics. **d.** Generalization performance on the extrapolation DMS stability task, where the test set is selected from the most peripheral proteins in the set (e, red), measured by In-Domain and transfer ratios for test set (left) and validation set (right). We took the same DMS dataset as c) but instead of using the LOPO procedure, selected a single test set consisting of the most distant proteins in sequence space visualized with an MDS of the Levenshtein distance matrix (see e). This most closely represents the setup in a protein engineering task, where the engineer is exploring outwords in sequence space from local knowledge. Here we see the strongest performance of UniRep over baselines, suggesting that UniRep is well suited to this protein engineering formulation. **e.** MDS of the DMS stability dataset Levenshtein sequence matrix with the test set (red) and training set from the stability extrapolation task (d) indicated. Villin, shown in 1D earlier, is enlarged and bolded. **f.** PCA of the DMS stability datasets UniRep distance matrix on with the test set (red) and training set from the stability extrapolation task (d) indicated. Villin, shown in 1D earlier, is enlarged and bolded. Note the example point, villin, and other test set points move closer to the training data in representation space compared to sequence space. **g.** PCA of the avGFP training set (blue) and FPbase test set (red) from Figure 4 with the best performing Doc2Vec model distance matrix. **h.** PCA of the avGFP training set (blue) and FPbase test set (red) from Figure 4 with the best performing UniRep model distance matrix. Note UniRep training data extends over the region of the test set, facilitating prediction by reducing extrapolation burden (which can be defined as the distance from each test set member to the nearest training observation).

**Supplemental Figure 11.**
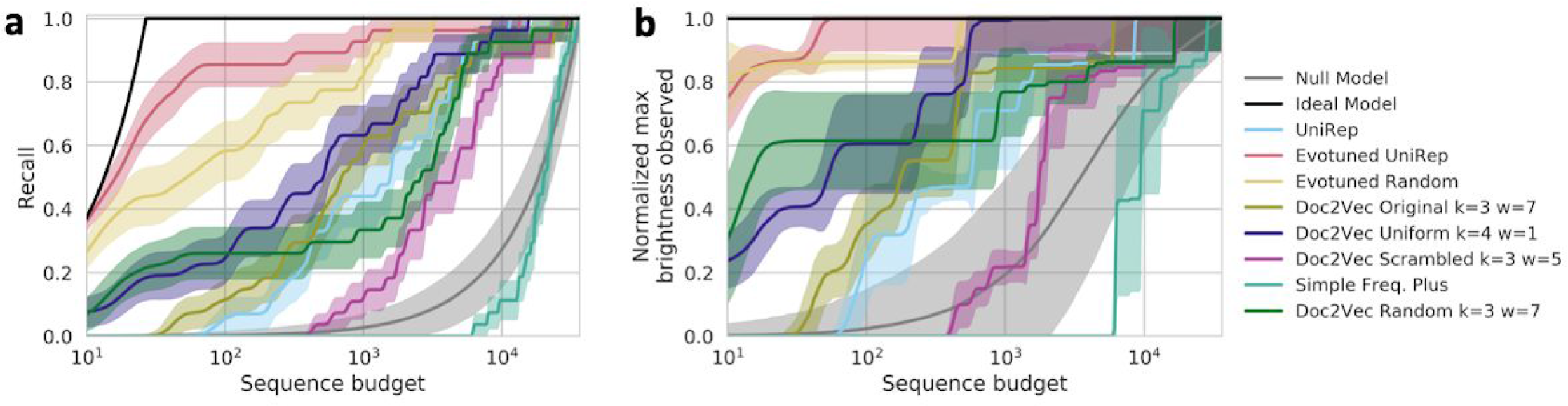
Efficiency curves for all baseline representations: recall (a) and maximum brightness (b). Note the variable performance of the best 4 variants of Doc2Vec presented in Yang *et al.* (2018) ^21^

**Supplemental Figure 12.**
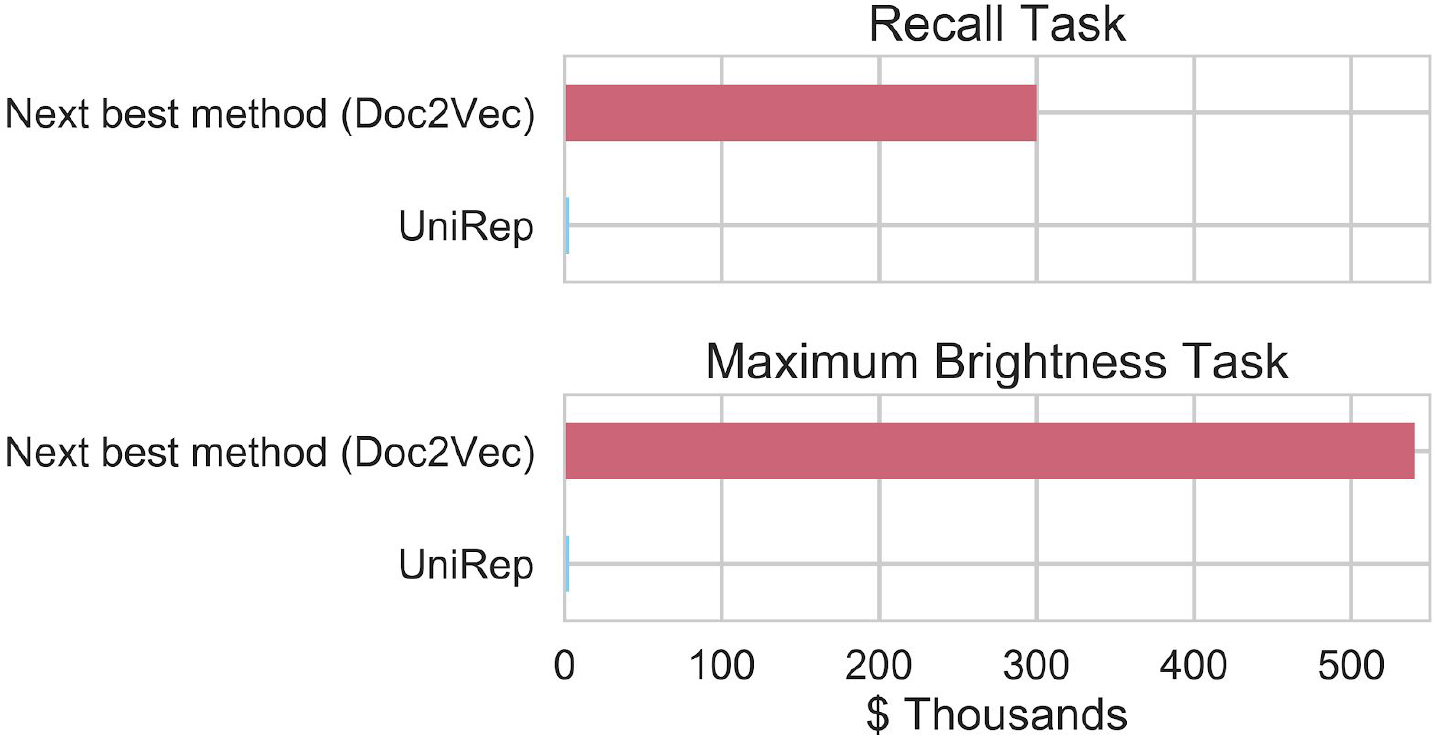
Estimated UniRep cost savings in GFP protein engineering tasks. Although real cost savings will vary depending on the number of non-functional sequences to sift through, Evotuned UniRep achieves 80% recall at ∼60 sequences tested, or approximately $3,000 assuming the most competitive full gene synthesis price of $0.07/nt^79^. By contrast, to achieve the same level of recall, the best Doc2Vec baseline would require ∼$300,000 (100x more). Random sampling still commonly used in this context would require $1,848,000 to achieve the same. Similarly, Evotuned UniRep captures the brightest sequence in the generalization set within the first $3,000 spent in testing, and the best Doc2Vec baseline would require ∼$540,000 (180x more) to do the same. Assuming on-target assembly rates improve and full economies of scale, multiplex gene assembly methods such as DropSynth^47^ could bring the cost of synthesizing a model proposed GFP down to ∼$2. At these cost rates, Evotuned Unirep would enable high purity functional diversity capture and function optimization for just a few hundred dollars. Taken together, these results suggest that Evotuning UniRep enables generalization to distant parts of the fitness landscape and thereby facilitates protein engineering by drastically minimizing the cost required to capture functional diversity and optimize function.

**Supplemental Table 6.** Analysis datasets and tasks.

| <u>Group</u> | <u>Protein(s) in the task</u> | <u>Size</u> | <u>Characteristic</u> | <u>Ref</u> |
| --- | --- | --- | --- | --- |
| Small-scale protein characteristics prediction | Cytochrome P450 | 261 | Thermostability | 21 |
|  | Bacterial Rhodopsin | 81 | Peak Absorption Wavelength |  |
|  | Epoxide Hydrolase | 152 | Enantioselectivity |  |
|  | Channelrhodopsin | 248 | Plasma Membrane Localization |  |
| Large-scale function prediction | TEM1b-lactamase | 5198 | Function (diverse, see Fig. 3e) | 35 |
|  | Ubiquitin - E1 activity | 1085 |  |  |
|  | Protein G (IgG domain) | 1045 |  |  |
|  | Pab1 (RRM domain) | 1188 |  |  |
|  | Ubiquitin | 1195 |  |  |
|  | PSD95 (Pd3 domain) | 1577 |  |  |
|  | Yap65 (WW domain) | 363 |  |  |
|  | Aminoglycoside kinase | 4234 |  |  |
|  | Hsp90 | 4021 |  |  |
| Large-scale stability prediction | hYAP65 | 829 | Stability (DMS data) | 5 |
|  | HHH_0142 | 775 |  |  |
|  | EEHEE_rd3_1716 | 775 |  |  |
|  | HEEH_rd3_0726 | 775 |  |  |
|  | EEHEE_rd3_1498 | 775 |  |  |
|  | EEHEE_rd3_1702 | 775 |  |  |
|  | HHH_rd2_0134 | 775 |  |  |
|  | HHH_rd3_0138 | 775 |  |  |
|  | HEEH_rd3_0872 | 775 |  |  |
|  | HEEH_rd2_0779 | 775 |  |  |
|  | HEEH_rd3_0223 | 775 |  |  |
|  | EEHEE_rd3_0037 | 775 |  |  |
|  | EHEE_rd3_0015 | 721 |  |  |
|  | EHEE_rd2_0005 | 721 |  |  |
|  | EHEE_0882 | 721 |  |  |
|  | Pin1 | 703 |  |  |
|  | villin | 631 |  |  |
|  | *De Novo* Proteins from Design Rounds | 56083 | Stability |  |
| GFP engineering | avGFP | 51715 | Brightness | 37 |
|  | A number of green fluorescent proteins | 27 |  | 39 |

**Supplemental Table 6.** Analysis datasets and tasks.
| Dataset | Size | Ref | Comment |
| --- | --- | --- | --- |
| SCOP 1.67 Superfamily Remote Homology Detection | 3802 | 62 | Representing 102 superfamilies |
| SCOP 1.67 Fold-level Similarity Detection | 3736 |  | Representing 86 folds |
| OXBench Database Reference Alignment | 811 | 26 | Classified into 180 families |
| Protein Family Prediction from HOMSTRAD Database | 3450 | 25 | Classified into 1031 families |

**Supplemental Figure 13.**
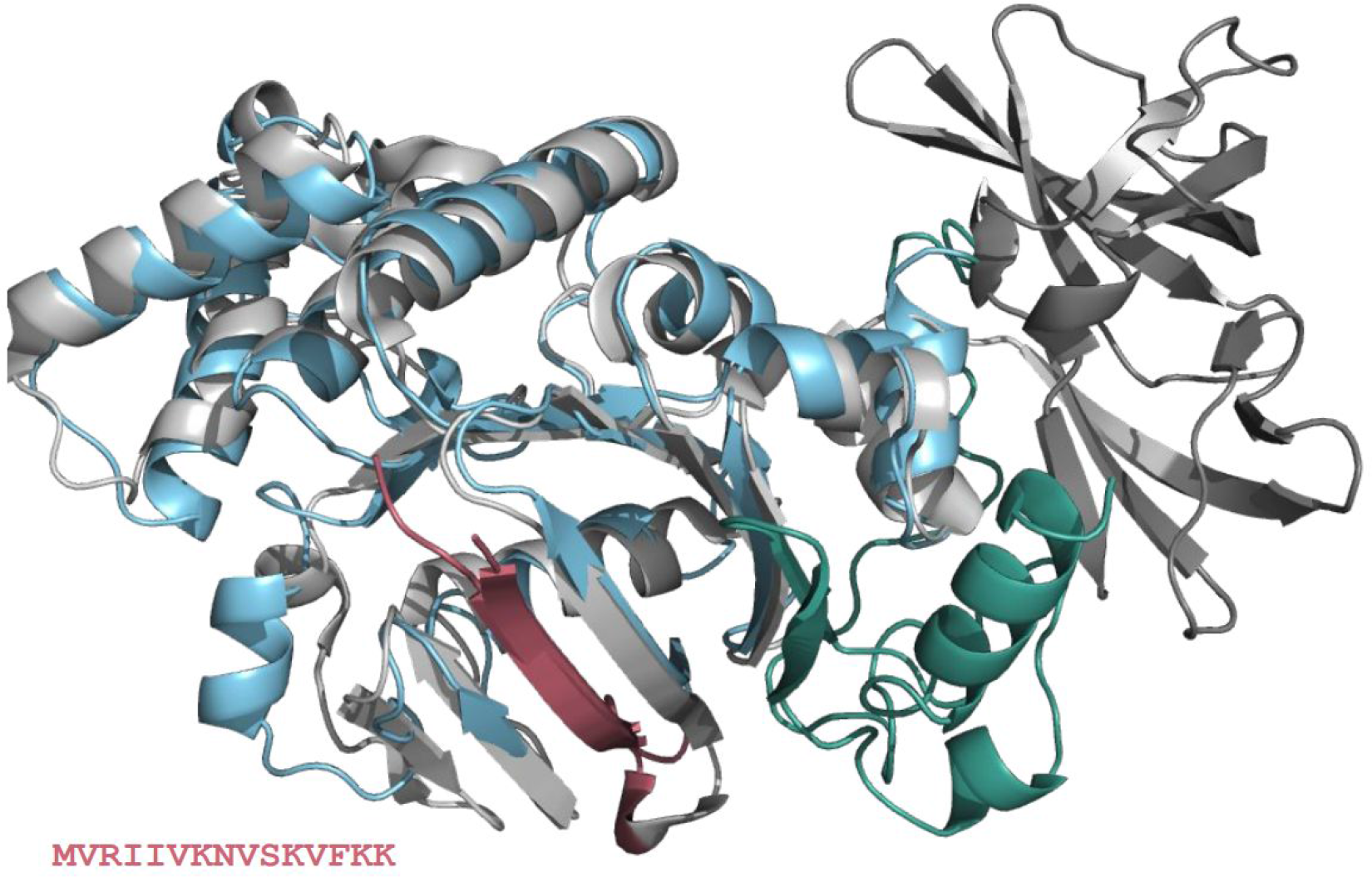
UniRep is a generative model of protein sequences. Homology model of a UniRep “babbled” sequence using a 15 amino acid seed (red) from a glucose ABC transporter (sequence from PDB:1oxx). Seed reference structure (PDB:1oxx) in grey. Modeled structure of babbled sequence shown in red (seed residues), blue (remaining N-terminus), and green (C-terminus which blasts to dipeptide ABC transporter with >40% identity). Alignment with the seed reference shows that UniRep has generated a sample which reconstructs structural regularities of the protein family. Full sequence blasts to ABC transporter family members with >50% similarity.

**Supp. Table 7.** Data augmentation of UniRep.

| Data Source | Motivation | Applications | Mechanism | Refs |
| --- | --- | --- | --- | --- |
| PDB crystal structures | Improve ab-initio structure prediction. | In-silico crystallography, structural comparisons | Joint training with the RGN, predicting next amino acid and structure to update joint weights by minimizing a hybrid averaged loss. | 16,52 |
| position sensitive scoring matrices (PSSMs) | Enrich representation of evolutionary relationships. | “Deep Homology”, semantic sequence comparisons, sequence search, phylogenomics. | Additional representation training input (predict next column of PSSM). |  |
| experimentally characterized synthetic mutants | Tune for a specific engineering task by training on synthetic sequences which have been synthesized and tested. | High-throughput model guided protein engineering. | Evotuning, but replace extant sequences with synthetic sequences which have measured bright. |  |
| ancestral sequence reconstruction | Tune for an engineering task by enriching training data with ancestral sequences near the target which have been shown to have desirable biotechnological properties. | Data-constrained model-guided protein engineering. | Evotuning, but replace extant sequences with high-confidence sequences from ancestral protein reconstruction methods applied to the target and nearby extant relatives. | 80 |
| Functional annotations (eg Pfam, GO, etc) | Incorporate existing functional data to improve representations for semantic sequence | Fast, vector math parallelized sequence annotation and homolog | Auxiliary tasks requiring the network to predict annotations after finishing predicting | 18,52 |
|  | search. | identification. | next amino acid up to the end of a sequence. |  |
| Experimental epistasis data | Train UniRep to capture higher-order features of protein fitness landscapes to model the relationship between micro and macro scale and the accordance with theoretical results. | Study of theoretical and experimental protein evolution. | Evotuning to the context of a target epistasis model then subsequent top model training on evotuned representations to predict experimental fitness values (as in Fig. 4) | 41,81 |

**Supp. Table 8.**
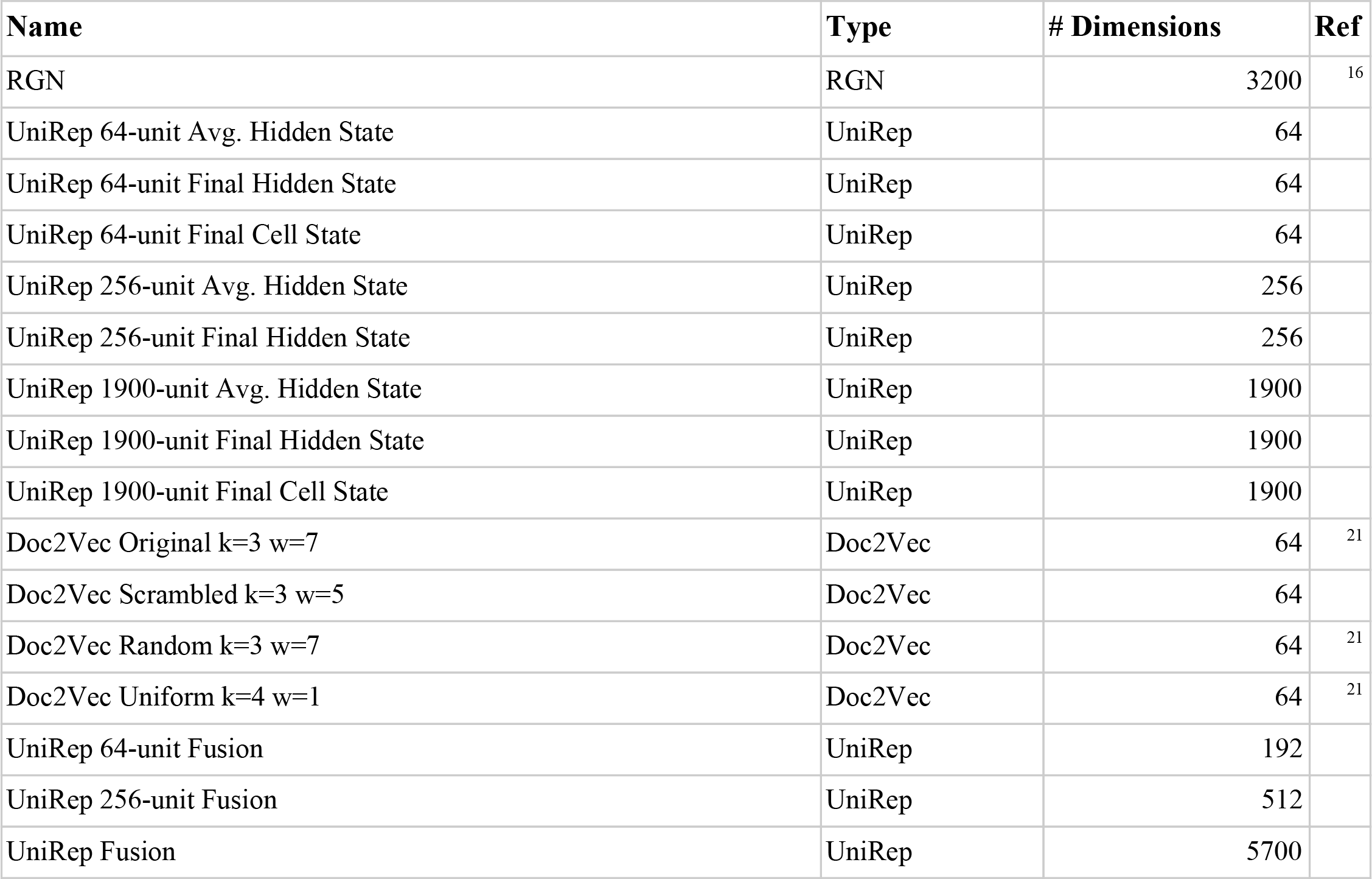
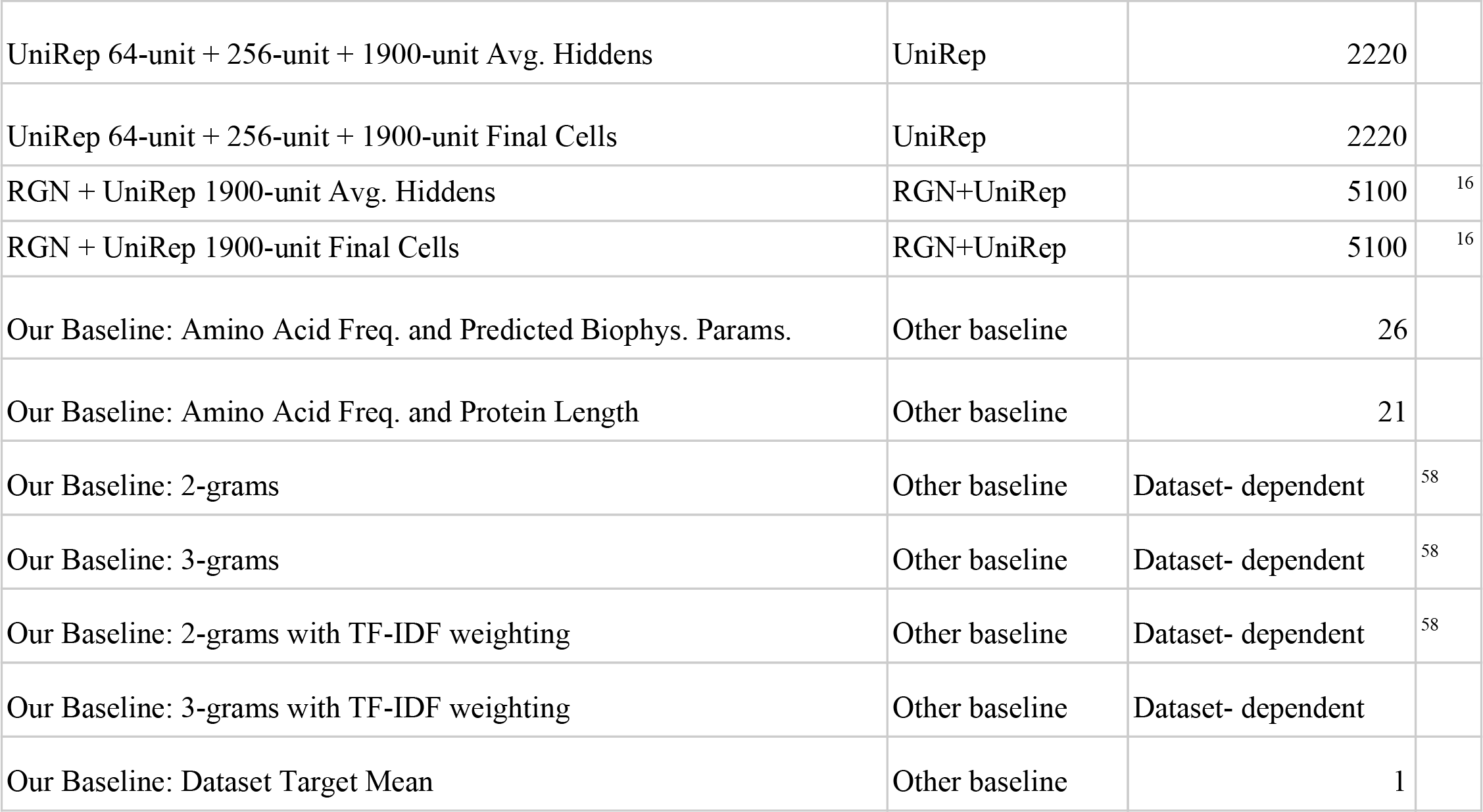
All the models we evaluated, including the baseline suite used for the majority of analyses in the manuscript. We additionally used Levenshtein distance (Needleman-Wunsch where all penalties are equal) for analysis in Fig. 2d and Rosetta total energy and NPSA measures for Fig. 3a (as described in Methods under “Stability Ranking Task”)

